# PKU tags, novel genetically encoded shape tags for cell labeling in light and electron microscopy

**DOI:** 10.1101/2025.11.25.690457

**Authors:** Rongbo Sun, Peilin Yang, Luyao Wang, Yixin Hu, Yini Yang, Fei Deng, Shaochuang Li, Mengyu Fan, Xiju Xia, Yulong Li

## Abstract

Distinguishing among neuronal cell types is crucial for deciphering complex neural networks and brain functions. However, the current repertoire of cell-labeling tools compatible with light microscopy (LM) and/or electron microscopy (EM) is limited compared to the vast number of cell types in the brain. Here, we introduce PKU (polymer king-size unit) tags, genetically encoded “shape tags” that leverage the polymerization of self-assembling proteins, spectrally distinct fluorescent proteins and, optionally, a nuclear targeting sequence to generate a series of multi-shaped (spherical or filamentous), multi-colored (blue, green, red, near-infrared) and multi-localized (cytosolic or nuclear) tags. By co-expressing multiple PKU tags within the same cell, a combinatorial strategy further expands the repertoire, which can theoretically yield hundreds of unique labeling patterns. Expressing PKU tags *in vivo* provides multi-cell-type labeling and neuronal circuit tracing, without altering the animal’s behavior or transcriptomic profiles. Moreover, when fused to the peroxidase APEX2, PKU tags maintain their shape-specific features, providing shape “barcoding” using EM. Thus, PKU tags represent a versatile and efficient toolkit for studying connectomics using both LM and EM.

## Main

The human brain contains thousands of specific neuronal cell types, each with distinct morphological, electrophysiological and transcriptional profiles^1^. To study these numerous cell types, understand their connectomic characteristics, and examine their respective roles in complex animal behaviors, a sufficient number of specific cell-labeling tags is needed for use in both light microscopy (LM) and electron microscopy (EM).

Currently, available optical imaging methods for labeling and identifying specific neuronal cell types rely primarily on 4–5 fluorescent proteins, each of which has distinct spectral properties. Brainbow and related cell-labeling strategies have demonstrated the ability to generate multiple fluorescent tags via the stochastic expression of multicolor fluorescent proteins within the same cell^2–8^. However, the overlapping spectra between fluorophores limit the application of this strategy, particularly when fluorescent indicators such as calcium sensors are also used. Although linear unmixing has been proposed as a method to separate highly overlapping fluorophores^9^, it requires careful selection of the fluorescent proteins, complex sample preparation and extensive post-analysis. Moreover, its effectiveness is often affected by the imaging conditions and background noise. Consequently, tags based solely on color are not sufficient. Attempts have been made to localize fluorescent proteins to specific cellular structures in order to expand the tag library^10^; however, the number of such tags is still remarkably low compared to the vast number of neuronal cell types, thereby underscoring the need for a much larger number of tags, particularly tags with orthogonal dimensions.

Compared to optical imaging, labeling and differentiating among neuronal cell types using EM is even more challenging. Immunoelectron microscopy, which is typically used for this purpose, is limited to currently available specific antibodies and different-sized gold nanoparticles, allowing for the simultaneous labeling of only two cell types at most; in addition, this method does not preserve cellular structure well and has suboptimal antibody penetration^11–13^. To expand labeling capacity, two general categories of genetically encoded tags have been developed for use in EM. The first category uses oxidized diaminobenzidine (DAB) to generate localized osmiophilic precipitates, providing multiple EM “barcodes” for subcellular labeling^14–19^; however, tag diversity is still limited, and identifying specific electron-dense labeling in lipid-rich membrane structures can be challenging. The second category includes metal-binding EM tags such as metallothionein-based tags^20–25^ and ferritin-like iron-sequestering nanoparticles^26–28^. Metallothionein-based tags require incubating live cells with toxic metals and have a laborious workflow; in addition, they provide suboptimal contrast against the general staining of cell structures, making them difficult to be distinguished and poorly suited to discern subtle differences in tag sizes. Ferritin-like iron-sequestering tags require culturing cells in an iron-rich medium and co-expressing transferrin and cargo proteins to enhance iron transport and sequestration; however, they still provide suboptimal electron density^27, 28^. Given these constraints, more effective high-contrast cell-labeling tags suitable for use in both LM and EM are clearly needed.

In nature, many proteins function as highly ordered, self-assembled polymers that form structured shapes such as fibers, rings, tubes, knots, and cages^29^. This wide structural diversity among protein complexes promoted us to examine whether these various shapes can be used to barcode distinct cell types **(Fig. 1a)**. We therefore screened 25 self-assembling proteins expressed in HeLa cells and found that 22 of these proteins successfully formed polymers, with distinct spherical and filamentous shapes. Given their composition of tens—or even hundreds—of monomers and their easily observable large sizes, we term these self-assembling monomers PKUs (polymer king-size units).

**Fig. 1.**
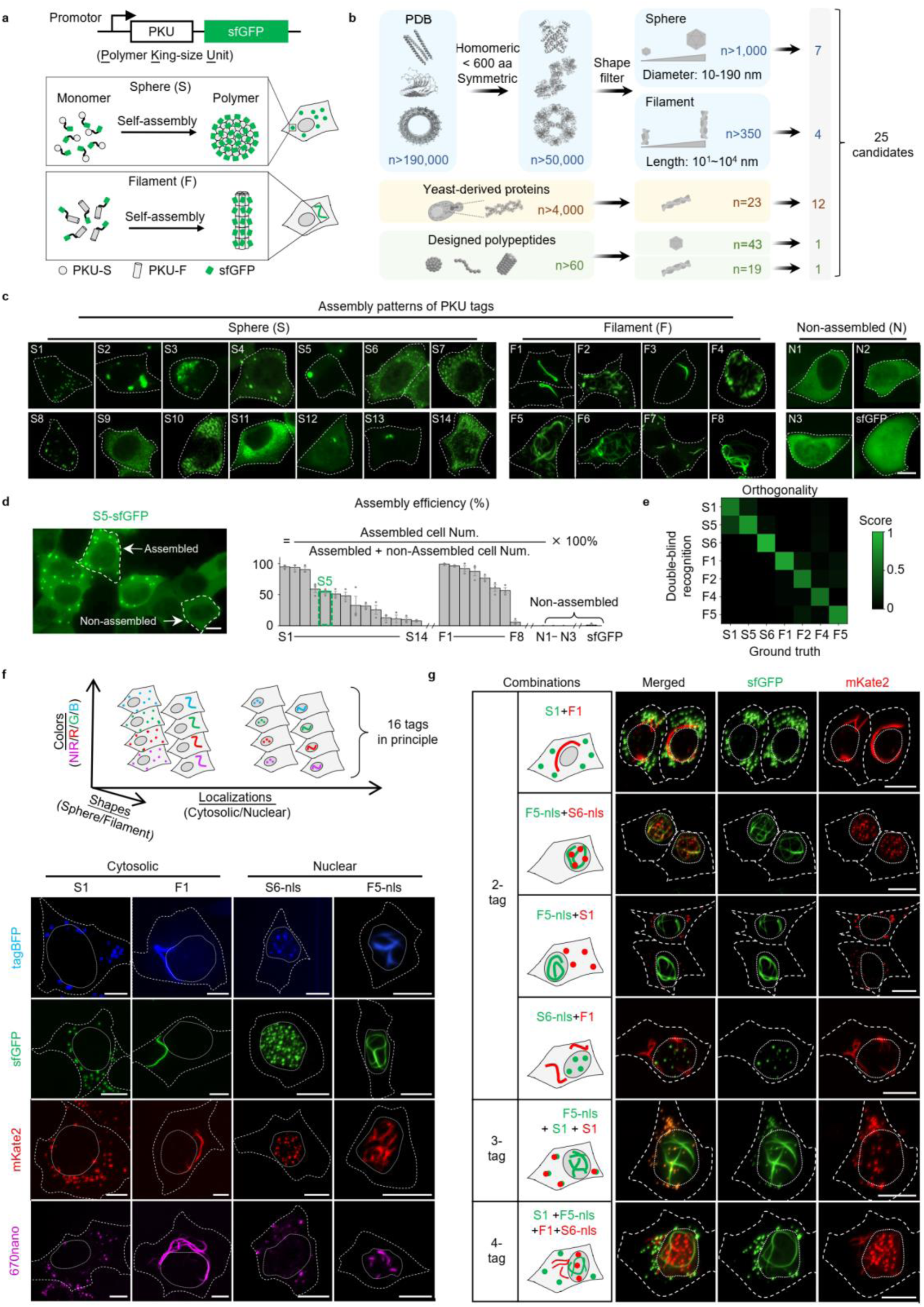
| Design and development of multi-shaped PKU tags. **a.** Schematic diagram depicting the principle of genetically encoded multi-shaped PKU tags, in which self-assembling protein monomers fused to sfGFP assemble to form polymers with specific shapes when expressed in cells. **b.** Sources used to identify PKU tag candidates. A total of 25 candidates were selected from the Protein Data Bank (PDB; 11 candidates), human homologs of self-assembling proteins derived from yeast (12 candidates), and *de novo* designed polypeptides (two candidates) based on reports in the literature. **c.** Assembly patterns of all 25 PKU tag candidates expressed in cultured HeLa cells, as well as control cells expressing sfGFP (bottom row, far-right). The PKU tag candidates are grouped into spherical tags (S1 through S14), filamentous tags (F1 through F8), and candidates that failed form a distinct shape (N1 through N3), and then sorted and ranked according to their assembly efficiency, from the highest (S1 and F1) to lowest (S14 and F8). In this and subsequent figures, the dashed lines indicate the cell outline. **d.** Left, expression of S5-sfGFP in HeLa cells, showing examples of cells with assembled patterns and non-assembled patterns; Right, quantification of the assembly efficiency of all 25 PKU-sfGFP tags (n> 37 cells from 3 coverslips). **e.** Heatmap showing double-blind recognition of three spherical and four filamentous PKU tags. **f.** Schematic diagram (top) and representative images (bottom) of 16 multi-dimensional labeling tags generated by a combination of two shapes (sphere or filament), four fluorescent colors (blue, green, red or near-infrared) and two subcellular localizations (cytosol or nucleus). Fifteen of the sixteen tags appear as expected, with the sole exception of S6-miRFP670nano-nls, which failed to localize to the nucleus. In this and subsequent figures, the dotted lines indicate the nucleus, based on DAPI or DRAQ5 staining. **g.** Combining multiple tags in the same cell generates new labeling patterns. Shown are examples in which two, three or four tags are expressed in the same HeLa cell. Scale bars: 10 μm.

We then fuse PKUs with different fluorescent proteins, with or without a nuclear targeting sequence, generating a series of multi-shaped, multi-colored, and multi-localized tags, each referred to as a PKU tag. Co-expression of multiple PKU tags within the same cell, as part of a combinatorial strategy, further expands the repertoire of possible labeling patterns. Moreover, PKU tags are suitable for *in vivo* labeling when expressed via adeno-associated virus (AAV) delivery or *in utero* electroporation (IUE), serving as reporters for use in both anterograde and retrograde neuronal circuit tracing. In addition, we developed multi-shaped PKU tags for use in EM, creating sphere-shaped and filament-shaped electron-dense labels. Thus, these PKU tags greatly expand upon the existing cell-labeling toolkit, providing a novel set of tools for examining the brain’s complexity using both LM and EM.

## Results

### Concept and development of multi-shaped PKU tags

To identify self-assembling proteins that can form the desired spherical or filamentous shapes in mammalian cells, we selected candidate proteins from three sources **(Fig. 1b)**. Firstly, we searched the Protein Data Bank (PDB), a vast repository of protein structures, and filtered for potential PKU candidates based on the following criteria: *i*) the proteins can form homomeric complexes, allowing for a simple expression system; *ii*) each monomer should be < 600 amino acids in size, allowing for AAV delivery; *iii*) the assembled complex exhibits symmetry, either through one or more axes of rotation forming defined, self-consistent assemblies (point group symmetry), or through a combination of rotation and translation that generates long, helical fibers (helical symmetry), both of which facilitate easy and precise shape recognition; and *iv*) relevant keywords such as “nanoparticle”, “filament”, “icosahedral”, and “octahedral” were used to focus on specific polymer shapes of interest. This filtering process reduced the entries to a few hundred candidates, from which 11 were randomly chosen. Secondly, we reviewed the literature and identified 23 yeast proteins that can form filamentous complexes^30, 31^, we then identified their human homologs and selected 12 candidates from this group. Lastly, we also included two *de novo* designed self-assembling proteins^32, 33^. Thus, a total of 25 candidates were chosen for multi-shape tag screening.

We then optimized the codons in these PKU candidates, fused superfolder GFP (sfGFP) to each candidate’s C-terminus via flexible linkers, and co-expressed each candidate with cytosolic RFP in HeLa cells. Among the 25 PKU candidates, 14 assembled to form spherical polymers (called S1–S14), 8 assembled to form filamentous polymers (called F1–F8), and three failed to show any obvious assembly patterns (called N1–N3) **(Fig. 1c)**. To compare assembly robustness of the various candidates, we quantified assembly efficiency by calculating the percentage of cells with assembled complexes **(Fig. 1d, Extended Data Fig. 1)**. The best-performing spherical and filamentous shaped candidates— S1 and F1, respectively—achieved assembly efficiency values up to 95% and 99%. We also found that the diameters of the spherical assemblies and the length of filament assemblies varied among the PKU tags, prompting us to determine whether assemblies within the same shape category can be distinguished reliably. We therefore conducted a double-blind test for three spherical and four filamentous PKU tags and found that spherical tags and filamentous tags were readily identified by subjects (with 99% accuracy) and specific PKU tags within the same shape category were also readily distinguishable (with 60%–94% accuracy) **(Fig. 1e)**.

We next examined whether the repertoire of PKU tags can be expanded further by combining them with other labeling strategies such as multicolor fluorescent proteins and subcellular localizations. We fused both S1 and F1 to four different fluorescent proteins (namely, tagBFP, sfGFP, mKate2, or miRFP670nano) and co-expressed each construct together with a spectrally separated cytosolic marker in HeLa cells, and found that both S1 and F1 retained their respective assembly shapes regardless of the fused fluorescent protein **(Fig. 1f)**. We also added a nuclear localization sequence (nls) to the C-terminus of the PKU-sfGFP tags with assembly efficiency in the top 50% (data not shown), and found that S6 and F5 successfully formed polymers in the nucleus while retaining their respective assembly patterns **(Fig. 1f)**; similarly, we expanded the nucleus-localized S6 and F5 tags by attaching four distinct fluorescent proteins **(Fig. 1f)**. Thus, in principle, by combining two shapes (spherical and filamentous), four colors and two subcellular localizations (cytosolic and nuclear), we could create a series of 16 distinct PKU tags. We found that 15 of these 16 PKU tags successfully assembled into their expected shapes, with correct subcellular localization; in contrast, S6-miRFP670nano-nls failed to localize to the nucleus **(Fig. 1f)**. In addition to using a single PKU tag for labeling, we attempted to combine several PKU tags in the same cell in order to generate new labeling patterns, thereby further expanding the tag library. We found that even when two, three, or four PKU tags were co-expressed in the same cell, each PKU tag assembled and was distinguishable based on its respective assembly pattern **(Fig. 1g**). By this combinatorial strategy, the tag library could, in principle, yield hundreds of unique labeling patterns according to combinatorial formulas. For example, choosing two tags from the 15 PKU tags yields 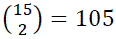 possible combinations; choosing three yields 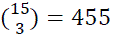 combinations; and choosing four yields 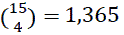 combinations. Moreover, we also demonstrated that expressing the PKU tags did not induce measurable cytotoxicity, even after 7 days **(Extended Data Fig. 2a-c)**.

### Labeling cultured neurons with multi-shaped PKU tags

Next, we tested the performance of PKU tags in cultured rat cortical neurons **(Fig. 2a)** by co-expressing the PKU-sfGFP tags with top 50% assembly efficiency (based on their expression in HeLa cells; data not shown) together with cytosolic RFP and found that all PKU tags tested retained their respective assembly capacity, with some tags showing labeling both in neuronal cell bodies and in neurites **(Fig. 2b)**. We also examined the aforementioned 16 orthogonal multiplex PKU tags and found that the majority formed their respective shapes, with the exception of three nls-fused tags that failed to localize to the nucleus, thus identifying a total of 13 PKU tags suitable for use in cultured neurons **(Fig. 2c)**. Further combinatorial expression strategies showed that co-expressing two or three tags in the same neuron produced clearly distinguishable patterns **(Fig. 2d)**.

**Fig. 2.**
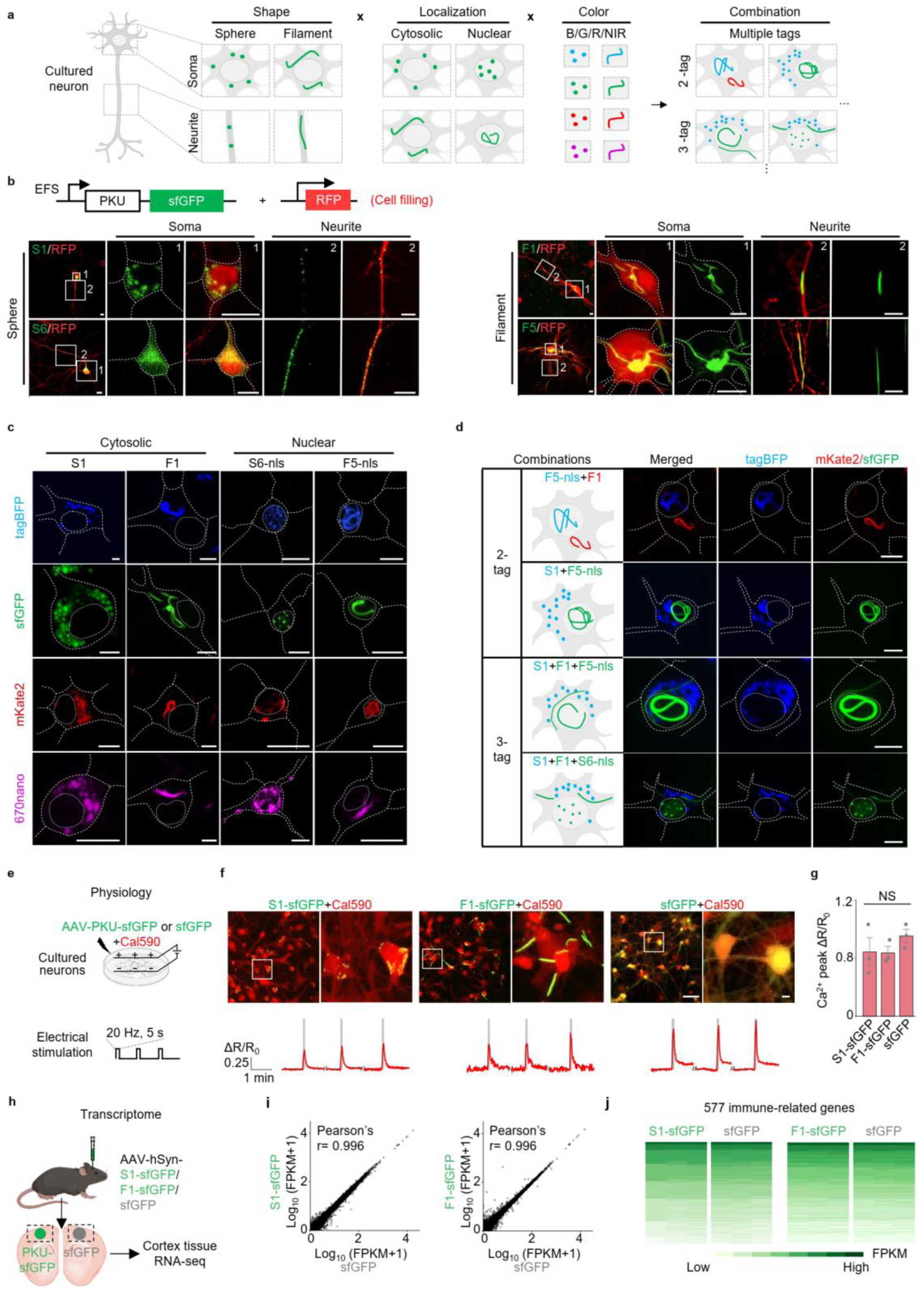
| Multi-shape PKU tag labeling in neurons does not affect cellular physiology or the transcriptome. **a**. Schematic diagram depicting the strategy used to label neurons with multiple PKU tags. **b.** Representative images of rat cortical neurons expressing spherical (left) and filamentous (right) PKU tags. PKU-sfGFP was co-expressed with a cytosolic red fluorescent protein. The white boxes indicate the magnified regions in the soma (1) and neurites (2). **c.** Representative images of cultured cortical neurons expressing 16 PKU tags generated by a combination of two shapes (sphere or filament), four fluorescent colors (blue, green, red, or near-infrared), and two subcellular localizations (cytosolic or nuclear). Thirteen of the sixteen tags express as expected; in contrast, S6-mKate2-nls, S6-miRFPnano-nls, and F5-miRFP670nano-nls failed to localize to the nucleus. **d.** Expressing multiple PKU tags in the same neuron generates distinct labeling patterns. The cartoons on the left depict the predicted assembly patterns. **e-g**. Expressing PKU tags has no measurable effects on cultured rat hippocampal neuron physiology. **e**. Schematic diagram depicting the experimental workflow, in which neurons expressing PKU-sfGFP or cytosolic sfGFP were loaded with Calbryte590 and electrically stimulated. **f**. Representative fluorescent images (top) and average calcium response traces (ΔR/R_0_) measured in neurons expressing S1-sfGFP (left), F1-sfGFP (middle), or cytosolic sfGFP (right). The red trace represents the mean ΔR/R_0_ among neurons from one coverslip (11–31 cells), while light gray traces represent responses measured in individual neurons. The dark gray shading indicates the electrical stimulation. **g**, Summary of peak ΔR/R_0_ measured in neurons expressing S1-sfGFP, F1-sfGFP or cytosolic sfGFP in response to electrical stimulation (n = 3 coverslips per group). Scale bars, 10 μm (**b**–**d** and **f**, right panels) and 100 μm (**f**, left panels) **h-j**. RNA-seq analysis confirms that expressing PKU tags does not alter the cellular transcriptome. **h**. Schematic diagram depicting the experimental strategy; PKU tag and sfGFP were expressed in the mouse motor cortex for 3 weeks, and the corresponding tissues were collected for RNA-seq. **i**. Pairwise comparisons of the transcriptomes between the motor cortex expressing S1-sfGFP vs. sfGFP (left) and between the motor cortex expressing F1-sfGFP vs. sfGFP (right); n=3 mice for each group. The Pearson′s correlation coefficients are indicated. **j**. The expression of selected immune-related genes was compared between motor cortex expressing PKU-sfGFP vs. sfGFP (n=3 mice per group).

To determine whether expressing the PKU tags affects neuronal physiology, we loaded neurons with calcium dye Calbryte590 and recorded calcium activity. We found no significant difference between PKU tag-expressing neurons and non-PKU tag-expressing neurons, indicating that the PKU tags had little effect on neuronal physiology **(Fig. 2e–g, Extended Data Fig. 2d–e)**. As an additional confirmation, we performed RNA sequencing (RNA-seq) analysis in mouse cortex samples expressing the PKU tags and found no significant effect on the cellular transcriptome **(Fig. 2h–j, Extended Data Fig. 2f-h)**.

### PKU tags can be used to label multiple cell types *in vivo*

To demonstrate further the feasibility of using PKU tags for multicellular labeling *in vivo*, we expressed four separate PKU tags in the mouse motor cortex via AAV injection or IUE **(Fig. 3a)**. We first expressed S1-tagBFP in excitatory neurons and found that the tag formed a clear spherical label **(Fig. 3b1)**, while Cre-dependent F1-mKate2 formed a clear filamentous label in the presence of Cre **(Fig. 3b2)**. We also expressed S6-sfGFP-nls and F5-tagBFP-nls, and found that—as expected—they formed spherical and filamentous complexes, respectively, in the nucleus **(Fig. 3b3–b4)**. Moreover, PKU tags retained their respective shapes for up to 9 weeks after virus injection **(Fig. 3e–g, Extended Data Fig. 3)**.

**Fig. 3.**
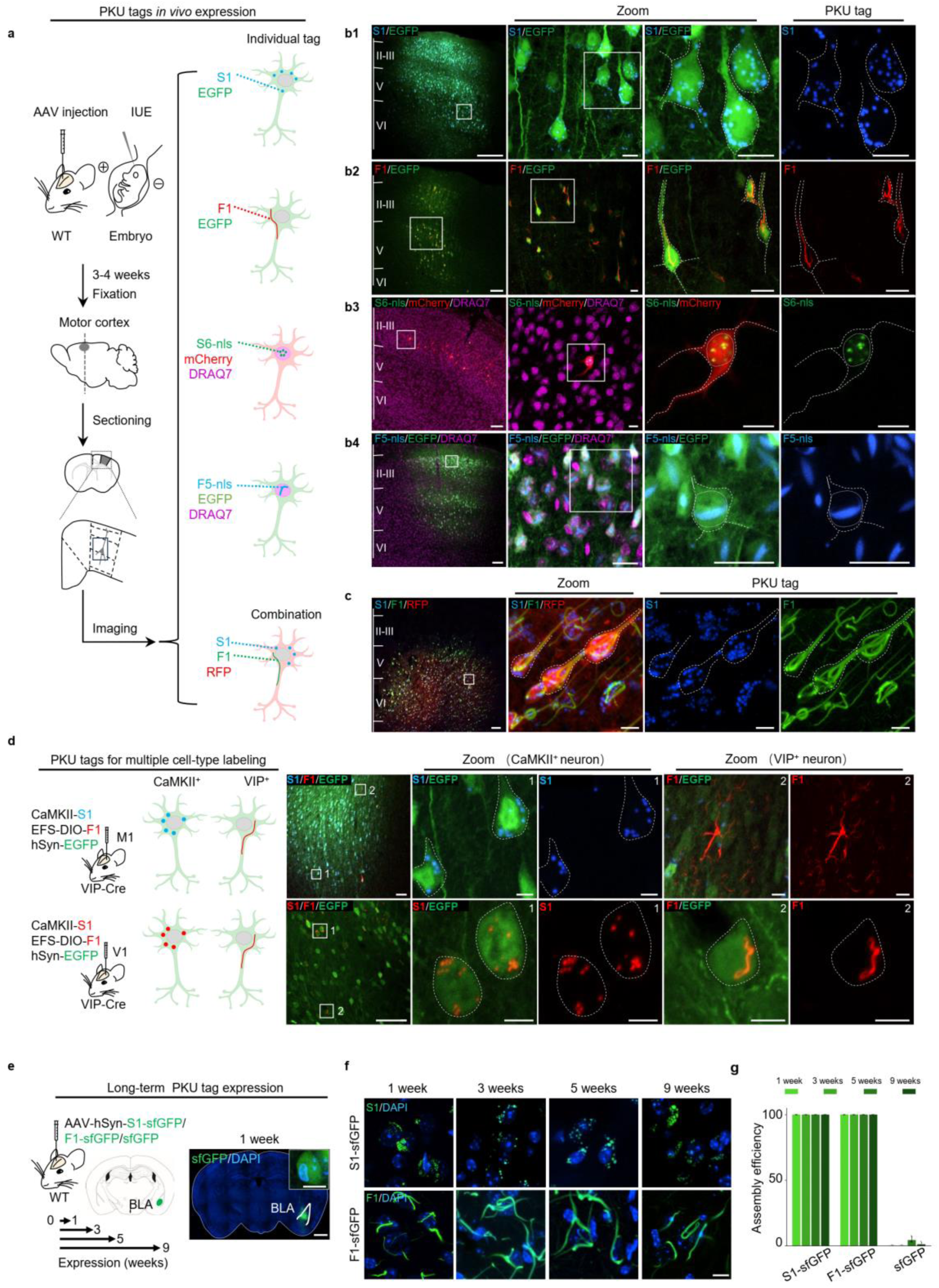
| PKU tags can be used for labeling multiple cell types *in vivo*. **a**. Schematic diagram depicting the experimental workflow in which PKU tags were expressed in the motor cortex by either AAV injection or *in utero* electroporation (IUE) at E14.5, followed by 3–4 weeks of expression, fixation, cryo-sectioning, and confocal imaging. Shown on the right are the predicted assembly patterns in **b1**–**b4** and **c**. **b.** Representative fluorescent images of sections of motor cortex expressing an individual PKU tag and a cytosolic marker. **b1**. AAV injection of CaMKII-S1-tagBFP and hSyn-EGFP. **b2**. AAV injection of EFS-DIO-F1-mKate2 and hSyn-Cre-P2A-EGFP. **b3**. IUE of EFS-S6-sfGFP-nls and CMV-mCherry, with nucleus staining using DRAQ7 (magenta). **b4**. AAV injection of CaMKII-F5-tagBFP-nls and hSyn-EGFP, with nuclear staining with DRAQ7 (magenta). The white boxes indicate increasingly magnified areas. Scale bars: 100 μm (first column) and 10 μm (second to fourth columns). **c.** Representative fluorescence images of sections of motor cortex co-expressing CaMKII-S1-tagBFP, hSyn-F1-sfGFP and hSyn-tdTomato. Scale bars same as **b**. **d.** Left, cartoon depicting the strategy for injecting virus mixture into VIP-Cre mice to label VIP^+^ inhibitory neurons and CaMKII^+^ excitatory neurons. Right, representative fluorescent images of two neuron types labeled with multi-shaped PKU tags either with the same color (top) or with distinct colors (bottom). Scale bars, 50 μm (first column), 5 μm (other columns). **e.** Left, schematic diagram depicting the strategy for expressing PKU tags in the mouse basolateral amygdala (BLA) for 1, 3, 5, and 9 weeks. Right, representative fluorescence image of a brain slice expressing sfGFP in the BLA; the nuclei were counterstained with DAPI (blue). The inset displays a magnified view of a single neuron. Scale bar: 1 mm, 10 μm (inset). **f.** Representative images of the indicated PKU tags 1, 3, 5, and 9 weeks after virus injection. Scale bars: 10 μm. **g.** Summary of the assembly efficiency of S1-sfGFP, F1-sfGFP and sfGFP measured at the indicated times after virus injection (n=3 brain slices per group).

To test whether multiple tags can be used *in vivo*, we co-expressed S1-tagBFP and F1-sfGFP in the motor cortex. As expected, each tag exhibited its respective shapes at the correct localization **(Fig. 3c)**. To test whether the PKU tags can be used to differentiate between different cell types, we expressed the S1 tag in excitatory neurons and expressed the F1 tag in VIP^+^ interneurons in VIP-Cre mice **(Fig. 3d)**. We found that the respective cell types could be labeled by specific shapes of PKU tags. Importantly, we found that mice expressing PKU tags for 3 weeks or 2 months in the basolateral amygdala (BLA)—a key brain region involved in fear and anxiety^34^—had no significant changes in fear-related and anxiety-related behavioral tests compared to control mice **(Extended Data Fig. 4)**.

### PKU tags can be used to trace neuronal circuit

Neurons generally receive their inputs from multiple upstream areas and send their output to multiple downstream regions. Commonly used strategies to investigate neuronal connectivity and brain function include anterograde and retrograde tracing^35,36^. Therefore, we tested whether PKU tags can be used for anterograde and/or retrograde tracing of neuronal circuits **(Fig. 4a)**.

**Fig. 4.**
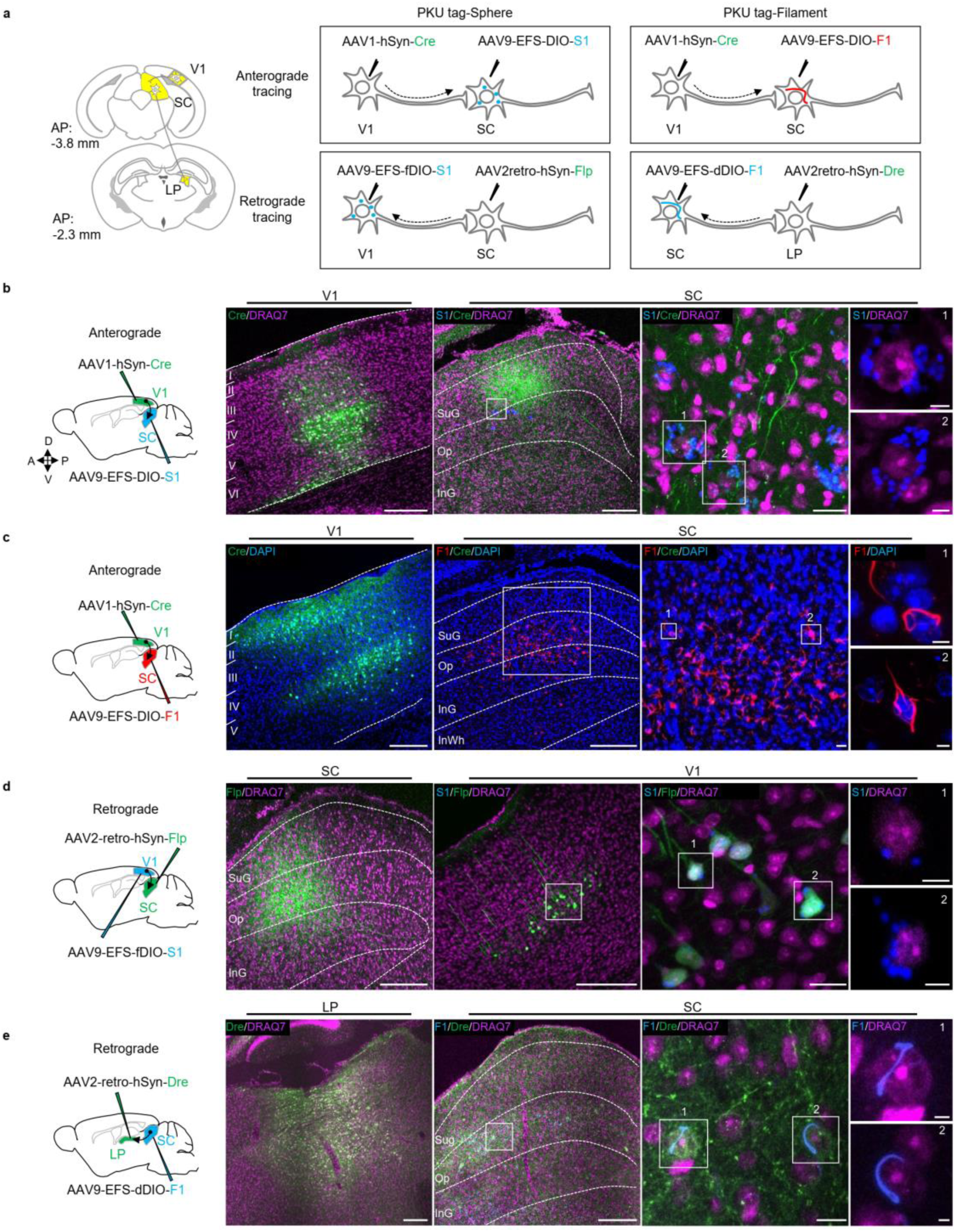
| PKU tags can be used to trace neuronal circuits. **a**. Experimental design for anterograde and retrograde neuronal tracing using PKU tags. Left, the anatomical locations and projections of visual area 1 (V1), the superior colliculus (SC), and the lateral posterior brain region (LP) are indicated. Right, the various PKU tags used to trace the indicated different circuits. **b.** Anterograde tracing of SC neurons receiving inputs from V1, using S1-tagBFP. Left, schematic diagram depicting the experimental design. AAV1-hSyn-Cre-P2A-GFP was injected into the V1 region of a wild-type mouse, followed by a second injection of AAV9-EFS-DIO-S1-tagBFP into the SC. Right, Cre-expressing neurons in the upstream V1 region (first panel) and S1-expressing neurons in the downstream SC region (second to fourth panels) induced by transsynaptic Cre recombinase. The nuclei were stained with DRAQ7 (magenta). **c.** Similar to **b**, except that AAV9-EFS-DIO-F1-mKate2 was injected, and the nuclei were stained with DAPI (blue). SC neurons that received inputs from V1 were labeled with red F1 tags. **d.** Retrograde labeling of V1 neurons that projecting to the SC, using S1-tagBFP. Left, AAV2-retro-hSyn-Flp-GFP was injected into the SC of a wild-type mouse, followed by a second injection of AAV9-EFS-fDIO-S1-tagBFP and AAV9-EFS-fDIO-EGFP (not shown) into V1. Right, Flp-expressing neurons in the SC (first panel) and S1-expressing neurons in the upstream V1 region (second to fourth panels) induced by retrograde expression of Cre from SC. **e.** Retrograde labeling of SC neurons projecting to the LP region, using F1-tagBFP. Left, AAV2-retro-hSyn-sfGFP-P2A-nls-Dre was injected into the LP region of a wild-type mouse, followed by a second injection of AAV9-EFS-dDIO-F1-tagBFP into the SC. Right, Dre-expressing neurons in the LP region and F1-expressing neurons in the upstream SC region induced by retrograde expression of Dre from the LP. Scale bars: 200 μm (first and second panels), 10 μm (third panel), and 5 μm (fourth panel).

The superior colliculus (SC) receives projections from visual area 1 (V1) in the cortex, and projects to the lateral posterior (LP) region^35^. We therefore expressed anterograde Cre-EGFP in V1, and either Cre-dependent S1 or F1 in SC **(Fig. 4b–c)**. In both cases, EGFP-positive cells were observed in V1, and the corresponding S1 or F1 tag was observed in the soma surrounding the nucleus in the downstream SC region, confirming anterograde tracing **(Fig. 4b–c)**; as a negative control, PKU tag expression in SC was absent when AAV9 serotype Cre recombinase was injected into V1 (data not shown). Next, we labeled upstream neurons in V1 that project to the SC region, as well as upstream neurons in the SC that project to the LP region **(Fig. 4d–e)**. We therefore expressed retrograde Flpo or Dre in the SC and LP regions, respectively, and Flpo-dependent or Dre-dependent S1 and F1 in the V1 and SC regions, respectively. In both cases, the appropriate PKU tags were observed in the upstream region, but was absent when retrograde recombinase injection was omitted (data not shown), thus confirming that PKU tags are also suitable for retrograde neuronal tracing.

### Screening and validation of PKU tags for multi-shape EM barcoding

The development of genetically encoded cell-labeling tags for EM is essential for studying the connectomics of complex nervous systems. However, the currently available tags are still limited. Therefore, we investigated whether our PKU tags can be adapted in order to generate a new, expanded series of EM barcodes. We therefore fused APEX2^15^ to the C-terminus of each validated self-assembled PKU tag **(Fig. 5a)** and assessed the assembly pattern in HeLa cells using DAB staining **(Fig. 5b)**. We found that all PKU-sfGFP-APEX2 tags retained their respective shapes, consistent with our fluorescence data **(Fig. 1c)**. We then quantified their assembly efficiency based on DAB imaging results and found that S1 and F1 again had the highest assembly efficiency **(Fig. 5c)** and were therefore used for our subsequent EM experiment. Our EM data revealed that cells expressing either S1-sfGFP-APEX2 or F1-sfGFP-APEX2 showed clearly distinguishable electron-dense labeling in a spherical or filamentous pattern, respectively; in contrast, non-transfected cells displayed no electron-dense structures **(Fig. 5d-e)**.

**Fig. 5.**
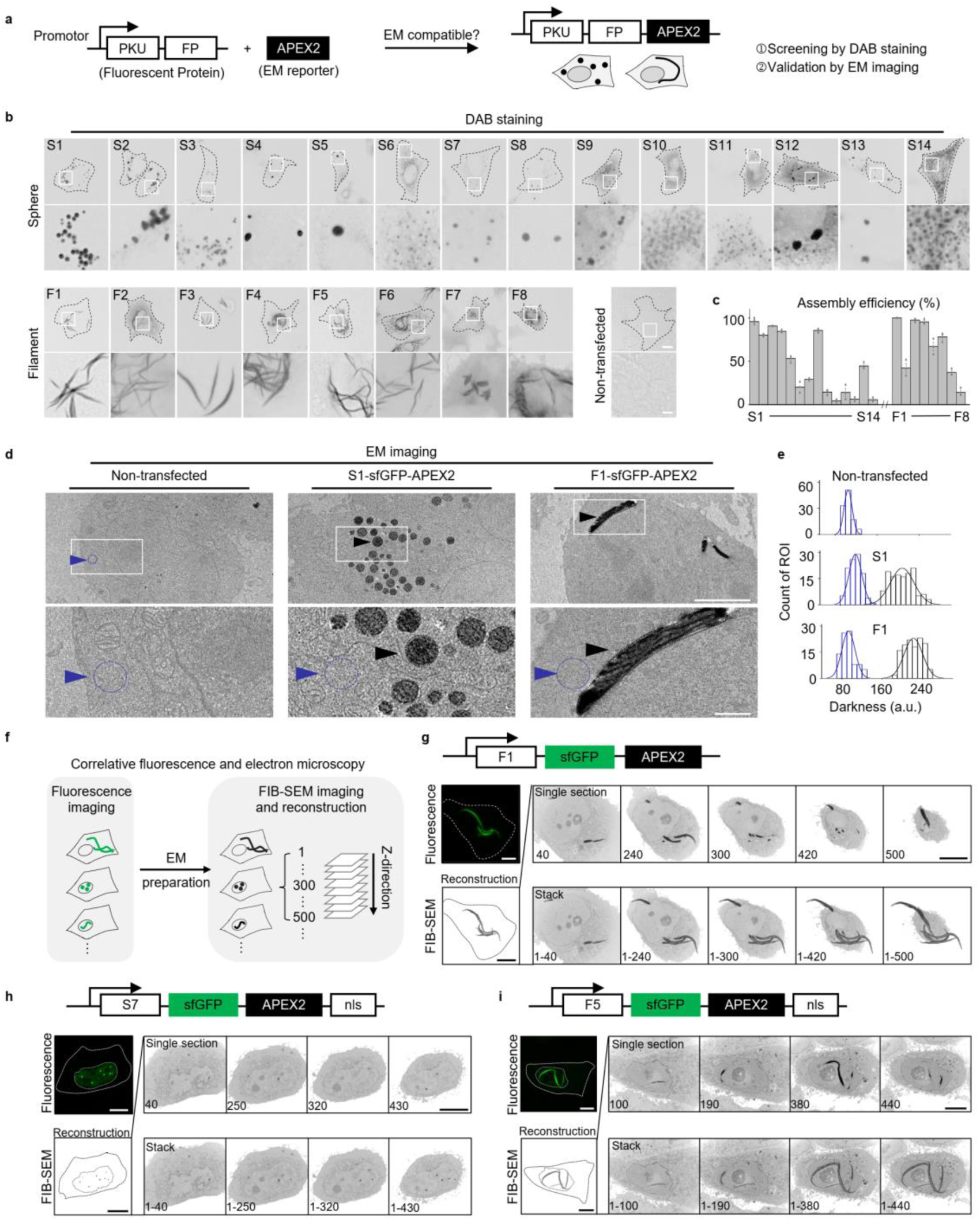
| Screening and validation of PKU tags for multi-shaped EM barcoding. **a**. Schematic diagram depicting the strategy for demonstrating that PKU tags are compatible with EM by fusing APEX2 to the C-terminus. **b.** Preliminary screening of all PKU-sfGFP-APEX2 fusion tags using DAB staining and brightfield microscopy. For each tag, the bottom image shows a magnified view of the region indicated by the white box in the top image. Scale bars: 20 μm (top) and 2 μm (bottom). **c.** Summary of the assembly efficiency of PKU-sfGFP-APEX2 fusion tags expressed in HeLa cells based on DAB staining results (n>200 cells from 3 coverslips). **d.** EM images of non-transfected control cells (left), cells expressing S1-sfGFP-APEX2 (middle) and F1-sfGFP-APEX2 (right). The top row displays whole-cell views, with white boxes indicating the regions shown in magnified views in the bottom row. The blue and red arrowheads highlight the areas used for quantifying the darkness in **e**. Scale bars: 5 μm (top) and 1 μm (bottom). **e.** Distribution of darkness in the cytosolic background (blue), S1-sfGFP-APEX2 (red), and F1-sfGFP-APEX2 (red). **f.** Schematic diagram illustrating CLEM (correlative fluorescence and electron microscopy). Cells expressing the PKU tags are first imaged using fluorescence microscopy, followed by EM preparation, ultra-thin sectioning (10-nm sections), and imaging using FIB-SEM (focused ion beam-scanning electron microscopy) and reconstruction. **g-i**. HeLa cells expressing F1-sfGFP-APEX2 (**g**), S7-sfGFP-APEX2-nls (**h**), or F5-sfGFP-APEX2-nls (**i**) were imaged by CLEM. Top left panels, maximum projection image. Bottom left panels, 3D reconstruction of intact electron-dense labels produced by PKU tags. Top right panels, EM snapshots of single sections, with black electron-dense labeling indicating the PKU tag signals. Bottom right panels, EM snapshots of single sections and stacked PKU tag signals. Dashed lines indicate cell outlines and dotted lines indicate the nucleus. Scale bars: 10 μm.

In a parallel comparison, we tested two previously reported spherical EM tags, 1M-Qt-nls and 2M-Qt, which were designed by fusing one or two copies of the heavy metal-binding protein metallothionein (M) with the self-assembling protein (Qt)^25^. These tags were shown to enable EM-based cell labeling in HEK cells, mouse brains and *Drosophila*. Following the published protocol, we observed some potentially distinguishable spherical electron-dense structures in the nuclei of cells expressing 1M-Qt-nls, but unfortunately not in the cytosol of cells expressing 2M-Qt, compared with non-transfected controls **(Extended Data Fig. 5)**. We quantified the darkness distribution of electron-dense labeling potentially generated by 1M-Qt-nls, along with the darkness of nuclear background regions (in Qt-expressing cells and untransfected cells) and other spherical-like structures (in untransfected cells). For cells expressing 2M-Qt, although we could not distinguish the electron-dense labeling potentially generated by 2M-Qt, we still quantified some dark labeling regions in the cytosol. The results showed that the darkness distribution of 1M-Qt-nls and 2M-Qt was slightly shifted toward higher values compared with background staining, but was comparable to that of other spherical-like EM structures. These results highlight the superior electron microscopy contrast provided by PKU tags.

Next, we performed correlative fluorescence and electron microscopy (CLEM) to test whether PKU tags can serve as a bridge between optical imaging and EM imaging **(Fig. 5f)**. We therefore expressed F1-sfGFP-APEX2 in cultured HeLa cells, imaged the sfGFP signal using LM, and then immediately prepared the samples for EM. We performed focused ion beam-scanning electron microscopy (FIB-SEM) in order to image nearly the entire cell and then reconstructed all of the F1-derived electron-dense signals observed in sequential thin sections, providing a complete set of F1 electron-dense labels that correlated well with the LM images **(Fig. 5g)**. In addition, we tested whether nucleus-localized PKU tags can serve as EM barcodes and found that both S7-sfGFP-APEX2-nls **(Fig. 5h)** and F5-sfGFP-APEX2-nls **(Fig. 5i)** produced the appropriately shaped, nucleus-localized electron-dense labels when expressed in HeLa cells.

### PKU tags can be used for multi-cell-type EM barcoding in neurons and astrocytes

Next, we examined whether our PKU tags can be used to label both mouse neurons and other cell types in culture and *in vivo*. For these experiments, we performed the following three steps: (1) fluorescence imaging to confirm successful expression and assembly of the PKU tags; (2) DAB staining and imaging to confirm APEX2 labeling; and (3) EM imaging **(Fig. 6a)**.

**Fig. 6.**
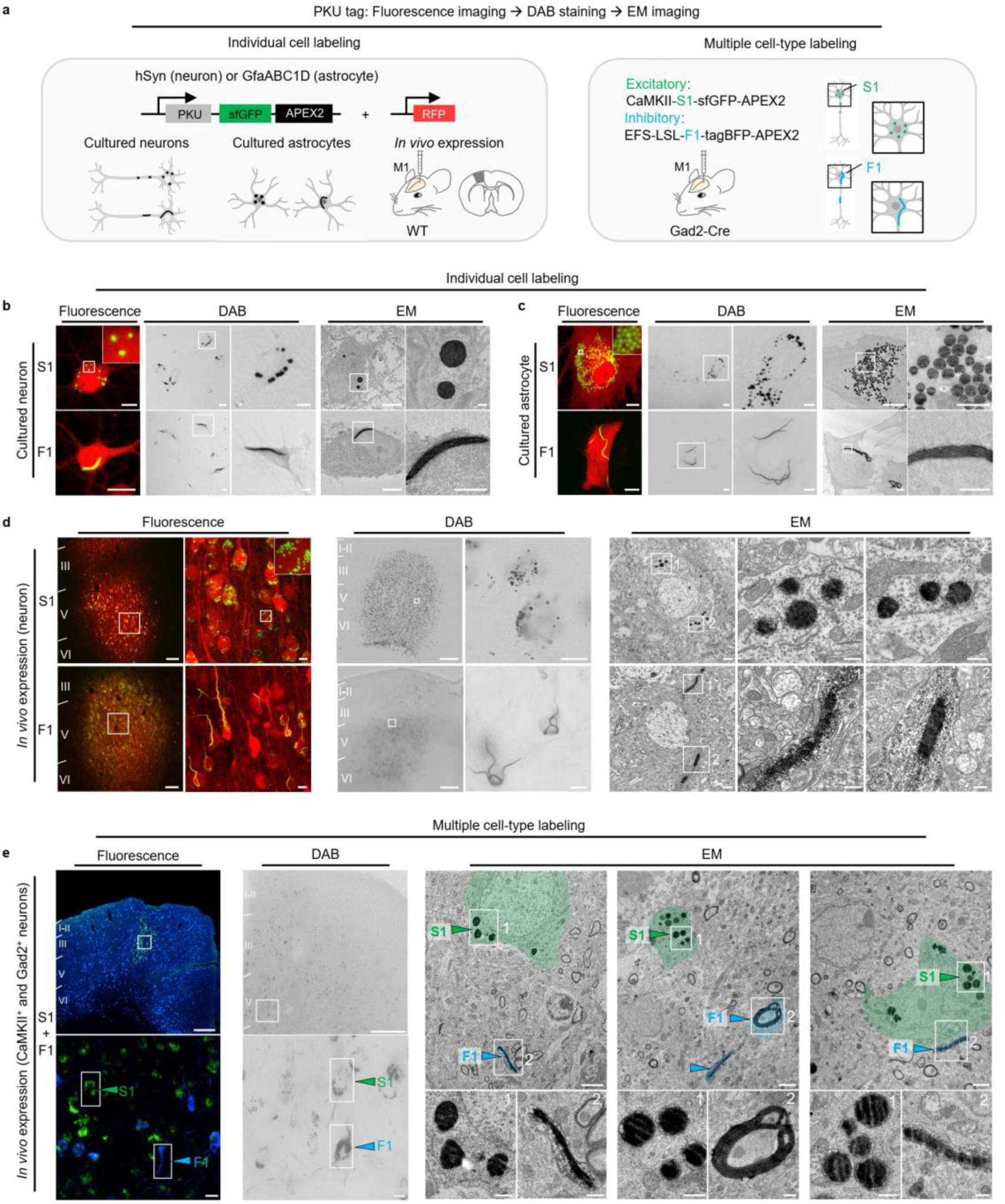
| PKU tags can be used for EM barcoding of neurons and astrocytes. **a**. Schematic diagram depicting the strategy for labeling an individual cell type in cultured neurons, cultured astrocytes, *in vivo* labeling in mice, and labeling multiple cell types in Gad2-Cre mice using PKU tags. **b.** Left columns, fluorescence images of neurons expressing S1-sfGFP-APEX2 (top) or F1-sfGFP-APEX2 (bottom). Middle columns, brightfield images of DAB-stained neurons expressing S1-sfGFP-APEX2 (top) or F1-sfGFP-APEX2 (bottom), with white boxes indicating regions magnified on the right. Right columns, EM images of electron-dense labeled neurons expressing S1-sfGFP-APEX2 (top) and F1-sfGFP-APEX2 (bottom), with white boxes indicating regions magnified on the right. Scale bars, fluorescence images, 10 μm (left); DAB images, 10 μm (left) and 5 μm (right); EM images, 5 μm (left) and 2 μm (right). **c.** Similar to **b**, except the experiments were conducted in cultured astrocytes. Scale bars same as **b**. **d.** Similar to **b**, except the experiments were conducted *in vivo* in mice. PKU tags were expressed in the motor cortex of the brain via virus injection. Scale bars: fluorescence images: 100 μm (left) and 10 μm (right); DAB images: 250 μm (left) and 10 μm (right); EM images: 2 μm (left), 0.5 μm (middle and right). **e.** Similar to **d**, except the excitatory neurons and inhibitory neurons in the motor cortex of Gad2-Cre mice were labeled with S1-sfGFP-APEX2 (green) and F1-tagBFP-APEX2 (blue), respectively. Left column, fluorescence microscopy images of the motor cortex, labeled with S1-sfGFP-APEX2 and F1-tagBFP-APEX2. Middle column, brightfield images showing DAB staining in the motor cortex of Gad2-Cre mice. In the left and middle columns, the boxes in the upper images indicate the areas magnified below and the boxes in the bottom images indicate the S1 and F1 labels. Right columns, EM images showing three fields of view of the motor cortex in Gad2-Cre mice expressing S1-sfGFP-APEX2 and F1-tagBFP-APEX2. In each field, the upper image shows an overview of both S1 and F1 labeling, while the bottom images show magnified views indicated by the white boxes in the upper image. Green and blue shading indicates S1-and F1-expressing neurons, respectively, while green and blue arrowheads indicate S1 and F1 labels, respectively. Scale bars: fluorescence images: 200 μm (top), 10 μm (bottom); DAB images: 200 μm (top), 10 μm (bottom); EM images: 2 μm (top) and 0.5 μm (bottom).

We first expressed the S1 and F1 tags in cultured neurons and astrocytes using cell type-specific promoters. Fluorescence imaging confirmed that S1 and F1 successfully assembled into the expected spherical and filamentous shapes, respectively, in both cell types, and this was confirmed by DAB staining and EM imaging **(Fig. 6b-c)**; in contrast, no labeling patterns were observed in non-transfected cells (data not shown). To test their feasibility *in vivo*, we expressed S1 and F1 separately in the motor cortex of mice via AAV injection and found robust assembly and labeling, with clear DAB deposits present only at the virus injection site and electron-dense labeling visible under EM **(Fig. 6d)**.

Lastly, we examined whether PKU tags could be used to simultaneously perform EM barcoding of different genetically defined neuronal populations. We therefore co-injected virus expressing CaMKII-S1-sfGFP-APEX2 and DIO-F1-tagBFP-APEX2 in the motor cortex of Gad2-Cre mice in order to simultaneously labeled excitatory and inhibitory neurons, respectively. Fluorescence imaging showed the appropriate shape labeling, corresponding to the appropriate cell types, with the fused fluorescent proteins serving as the ground truth **(Fig. 6e)**. After confirming successful labeling using DAB staining, we then prepared the samples for EM. The resulting EM images clearly showed spherical and filamentous electron-dense labeling, denoting the cell types. Thus, we showed that PKU tags can be used to label multiple cell types, underscoring their potential for use in ultrastructural analysis and neural circuit tracing, as well as to study connectomics using EM.

## Discussion

Here, we report the development and validation of a new series of genetically encoded PKU tags that integrate the polymerization of self-assembling proteins with spectrally distinct fluorescent proteins and, optionally, a nuclear targeting sequence, generating multi-shaped, multi-colored, and multi-localized labeling patterns. These PKU tags are likely to serve as a valuable tool for tracing neuronal circuits across multiple brain regions. Furthermore, to broaden the scope of multi-shape labeling using EM, we fused the PKU tags with APEX2, creating a new set of genetically encoded EM reporters that retained their shapes and provided clear electron-dense labels, even under highly stringent fixation conditions. In summary, our PKU tags provide a versatile, efficient toolbox to label cells for examination using both LM and EM.

Importantly, PKU tags provide several advantages over other tags, First, they are genetically encoded, allowing for expression in specific cell types. Second, the size of each monomer is compatible with AAV packaging, making this strategy suitable for use *in vivo*. Third, they provide a bright signal due to the presence of multiple copies of the fluorescent proteins in assembled complex, thereby reducing the laser intensity needed for imaging and reducing phototoxicity. Finally, they have no apparent effects on cell viability, cellular physiology, the transcriptome, or animal behavior.

With respect to their use in EM, PKU tags also have several advantages over existing tools. First, they provide clearer and significantly stronger electron-dense labeling due to the enzymatic amplification of multiple copies of APEX2, making the PKU tags much easier to distinguish from cellular structures. Second, the expression system is relatively simple, requiring the introduction of only a single protein, APEX2, unlike the more complex iron-sequestering EM methods that require co-expression of transferrin and cargo proteins to enhance iron transport and sequestration. Third, the EM protocol used in this study is both simple and robust, as it does not require culturing cells in an iron-rich medium or performing heavy-metal staining with lead, a procedure that can easily cause contamination, as observed in our experiments. Among the theoretical 16 tags, we successfully demonstrated 15 when expressed in HeLa cells and 13 when expressed in cultured neurons. Although not all 16 tags were realized, we believe this could be achieved by screening a broader range of self-assembling proteins. Notably, we found that high expression of nucleus-targeted PKU tags can lead to “leakage” of the tags outside of the nucleus; this effect should be minimized, particularly when the tag is used in combinatorial applications. Moreover, we found that S7-sfGFP-APEX2-nls monomers failed to assemble into spherical polymers in the nucleus in cultured neurons, while F5-sfGFP-APEX2-nls largely leaked into the cytoplasm, indicating that further optimization is needed before these tags can be used.

Beyond labeling specific cell types, we also demonstrated that PKU tags can be used for neuronal circuit tracing, a field where PKU tags may have significant potential for uncovering complex neuronal connections. For example, by using distinct PKU tags combined with multiple orthogonal recombinase systems and tracing tools, we could theoretically map the detailed connections of each neuron within a specific brain region to multiple upstream and downstream areas in the same mouse. If a neuron expresses multiple PKU tags due to its involvement in various projections, expansion microscopy can be employed to resolve distinct shapes of the PKU tags.

In addition, PKU tags may also have potential applications with respect to developing multi-shaped probes, enabling the simultaneous imaging of probe signals in multiple cell populations in a single channel. Pursuing a similar goal of dissecting detailed information in a single imaging session, Linghu et al. introduced the signaling reporter islands (SRI) method^37^, which fuses self-assembling peptides with multiple probes in order to form clusters that can simultaneously detect several molecular probes such as calcium, cAMP, and protein kinase A (PKA) within a single cell. However, this method is relatively limited, as the signaling molecules appear as indistinguishable puncta, requiring subsequent cell fixation and antibody staining in order to identify specific molecules. In theory, incorporating the concept of shape can increase the number of distinguishable labels and potentially allow for identification even in live cells without the need for fixation.

In summary, our novel PKU tags provide a promising new strategy for labeling neurons and other cell types. Thus, PKU tags will likely facilitate biological studies, particularly studies designed to unravel the complex network architecture of neural circuits by providing detailed EM-derived connectome data in genetically defined cell types.

## Acknowledgments

This work was supported by grants from the National Natural Science Foundation of China (31925017 to Y.L.), the National Key R&D Program of China (2022YFE0108700 and 2023YFE0207100 to Y.L.), the Beijing Municipal Science & Technology Commission (Z220009 to Y.L.), and the NIH BRAIN Initiative (1U01NS120824 to Y.L.). Support was also provided by the Feng Foundation of Biomedical Research, the New Cornerstone Science Foundation through the New Cornerstone Investigator Program, the Peking-Tsinghua Center for Life Sciences, the State Key Laboratory of Membrane Biology at Peking University School of Life Sciences (all to Y.L.).

We thank Yiqun Liu, Yingchun Hu, Jun Hu, Yingying Guo, and Xinpeng He at the School of Life Sciences at Peking University for their generous help regarding EM techniques. We thank Xiaoguang Lei at PKU-CLS, the optical Imaging platform, and the small animal imaging platform of the National Center for Protein Sciences at Peking University in Beijing, China, for their support and assistance with the Operetta CLS high-content imaging system, the Nikon A1RSi+ laser scanning microscope, and the animal behavior facility. We thank Professor Jilong Liu at ShanghaiTech University for providing the *mCtps1* gene. We also thank Professor Miao He at Fudan University for providing the VIP-Cre transgenic mice. Finally, we thank the members of the Li lab for helpful suggestions and comments on the manuscript.

## Author contributions

Y.L. supervised the project. R.S. developed and validated the PKU tags in cultured HeLa cells and neurons, with assistance from Y.H., Y.Y and P.Y. R.S. validated the performance of the PKU tags *in vivo* with assistance from L.W., S.L., P.Y., M.F., and X.X. L.W. performed the neuronal circuit tracing experiments. R.S. performed the EM experiments with assistance from P.Y. P.Y. performed the cell viability assay. R.S. performed the experiments related to physiological measurements and transcriptomic analysis. F.D. performed the behavior assays with assistance from R.S. and P.Y. All authors contributed to the interpretation and analysis of the data. R.S. and Y.L. wrote the manuscript with input from all authors.

## Methods

### Cell lines and culture conditions

HeLa cells were a generous gift from Yi Rao (Peking University). HEK293T cells were purchased from ATCC (CRL-3216). All cell lines were cultured in Dulbecco’s Modified Eagle’s Medium (DMEM; Biological Industries, 06-1055-57-1ACS) supplemented with 10% (v/v) fetal bovine serum (FBS; CellMax, SA301.02), and 1% (v/v) penicillin-streptomycin (Gibco, 15140122) at 37°C in humidified air containing 5% CO_2_.

### Primary cell cultures

Rat primary cortical and hippocampal neurons were obtained from postnatal day 0 (P0) Sprague-Dawley rat pups of both sexes (Beijing Vital River Laboratory Animal Technology Co., Ltd.). In brief, cortical and hippocampal neurons were dissociated from the dissected brain using 0.25% trypsin-EDTA (Gibco, 25200056), plated onto 12-mm glass coverslips precoated with poly-D-lysine hydrobromide (Sigma, P7280), and cultured in Neurobasal medium (Gibco, 21103049) containing 2% (v/v) B-27 supplement (Gibco, A3582801), 1% (v/v) GlutaMAX (Gibco, 15140122), and 1% (v/v) penicillin–streptomycin (Gibco, 15140122) at 37°C in humidified air containing 5% CO_2_.

Rat primary cortical astrocytes were also obtained from P0 Sprague-Dawley rat pups of both sexes. In brief, mixed cortical cells were isolated from the cerebral cortex using 0.25% trypsin-EDTA and plated on poly-D-lysine-coated T25 flasks. The culture medium contained DMEM supplemented with 10% (v/v) FBS and 1% (v/v) penicillin-streptomycin and was changed the day after plating, and every 2 days thereafter. After 7–8 days in culture (DIV7 or DIV8), the flasks were shaken at 180 rpm for 30 min to remove microglia, then at 240 rpm for 6 hrs to remove oligodendrocyte precursor cells. The remaining astrocytes were dissociated with trypsin-EDTA and plated on 12-mm glass coverslips in 24-well plates containing culture medium at 37°C in a humidified air containing 5% CO_2_.

### Mice

All procedures involving animals were performed using protocols approved by the Animal Care and Use Committee at Peking University. Adult female C57BL/6J mice were obtained from Beijing Vital River Laboratory Animal Technology Co., Ltd. VIP-Cre mice (JAX Strain 010908) were generously provided by Professor Miao He’s laboratory at Fudan University. Gad2-Cre mice (JAX Strain 019022) were obtained from the Jackson Laboratory. Mice (6–9 weeks of age) were used for the *in vivo* experiments. All mice were group-housed (up to five mice per cage) in a temperature-controlled (18–23°C) and humidity-controlled (40–60%) room with a 12-h/12-h light/dark cycle, with food and water available ad libitum.

### Molecular biology

All plasmids were constructed using the Gibson assembly method and verified by Sanger sequencing. cDNAs encoding the PKU tags were either PCR-amplified from the human cDNAs (hORFeome database 8.1) or mammalian-codon optimized and synthesized by Huada Qinglan Biological Technology. Mutations were introduced to reduce protease activity, including S1 (human PRDX4), S2 (human PRDX3), and F1 (mouse Ctps1). The mutation sites are as follows: S1: p.Cys124Ser and p.Cys245Ser; S2: p.Cys108Ser and p.Cys229Ser^38–40^; F1: p.Ser462Ala^41^. Nucleus-targeted PKU tags were fused with a nuclear localization signal (nls) at the C-terminus. The PKU tags expressed in cultured HeLa cells, HEK293T cells, and astrocytes were cloned into the pAAV vector under the control of the EF-1 alpha short (*EFS*) promoter or the astrocyte-specific *GfaABC1D* promoter. The APEX2-fused PKU tags expressed in cultured HeLa cells were cloned into the Pacific vectors under the control of the CAG promoter. The PKU tags expressed in cultured neurons and *in vivo* were cloned into the pAAV vector under the control of the human synapsin (*SYN1*) promoter (for pan-neurons), the *CaMKII* promoter (for expression in excitatory neurons), or the *EFS* promoter (for recombinase-dependent cases) for AAV packaging.

### AAV production

AAVs expressing PKU tags were packaged at WZ Biosciences. Other AAVs were purchased from BrainVTA (Wuhan, China), Braincase, or WZ Biosciences. The AAVs were used to infect cultured primary cells or were injected into specific brain regions for *in vivo* applications.

### DNA Transfection and AAV infection in cultured cells

For the experiments in **Fig. 1**, **Fig. 5** and **Extended Data Fig. 2b**, HeLa or HEK293T cells were plated in 24-well or 96-well plates and grown to 50–70% confluence before transfection with PEI (1 μg plasmid and 3 μg PEI per well in 24-well plates, or 300 ng plasmid and 900 ng PEI per well in 96-well plates). After 6–8 hrs, the medium was replaced with standard culture medium, and the cells were imaged 24−48 hrs or 7 days after transfection.

For the experiments in **Fig. 2b-c**, cultured neurons were transfected using the calcium phosphate method. In brief, DIV 6−8 neurons were cultured with a mixture containing 125 mM CaCl_2_, Hank’s Balanced Salts (HBS, pH 7.04–7.12), and 2 μg DNA. After incubation for 2 hrs, the DNA-Ca_3_(PO_4_)_2_ precipitate was removed by washing the coverslips with preheated (37°C) HBS (pH 6.80). Imaging was performed 3−5 days after transfection.

For the experiments in **Fig. 2d**, at DIV2 the cultured neurons were transduced with the following mixtures of viruses: AAV9-CaMKII-F5-tagBFP-nls, AAV9-EFS-DIO-F1-mKate2, and AAV9-hSyn-EGFP-P2A-Cre (first row); AAV9-CaMKII-S1-tagBFP, AAV9-hSyn-F5-sfGFP-nls, and CaMKII-mKate2 (second row); AAV9-CaMKII-S1-tagBFP, AAV9-hSyn-F1-sfGFP, AAV9-hSyn-F5-sfGFP-nls, and CaMKII-mKate2 (third row); and AAV9-CaMKII-S1-tagBFP, AAV9-hSyn-F1-sfGFP, AAV9-hSyn-S6-sfGFP-nls, and CaMKII-mKate2 (fourth row). The cells were imaged 6−8 days after infection.

For the experiments in **Fig. 2e**-**g** and **Extended Data Fig. 2d**-**e**, at DIV2 the cultured neurons were transduced with AAV9-hSyn-S1-sfGFP, AAV9-hSyn-F1-sfGFP, AAV9-hSyn-S6-sfGFP-nls, AAV9-hSyn-F5-sfGFP-nls, AAV9-hSyn-sfGFP, or AAV9-hSyn-sfGFP-nls. The cells were imaged 9 days after infection.

For the experiments in **Fig. 6b**, cultured neurons were transduced with AAV9-hSyn-S1-sfGFP-APEX2 or AAV9-hSyn-F1-sfGFP-APEX2 together with AAV9-CaMKII-mKate2.

For the experiments in **Fig. 6c**, cultured astrocytes were transfected with PEI (1 μg plasmid and 3 μg PEI per well in 24-well plates) and imaged 3 days later.

Where indicated, the nuclei were counterstained with DAPI (Beyotime), DRAQ5 (BioLegend), or DRAQ7 (BioLegend).

### Fluorescence imaging of cultured cells and primary cells

HeLa cells and cultured primary cells were imaged using an Operetta CLS high-content screening system (PerkinElmer) or a Ti-E A1R inverted confocal microscope (Nikon) at various *z*-planes, and maximum intensity projection was performed to generate the final images. The Operetta CLS system was equipped with a 40×/1.1 NA water-immersion objective, a 63×/1.15 NA water-immersion objective, a 390–420 nm LED laser, a 460–490 nm LED laser, a 530–560 nm LED laser, and a 650–675 nm LED laser, and was controlled using Harmony 4.9 software. The BFP, GFP, RFP and near-infrared fluorescent signals were collected using 430–500 nm, 525–580 nm, 570–650 nm, and 655–760 nm emission filters, respectively. The confocal microscope was equipped with a 10×/0.45 NA objective, a 20×/0.75 NA objective, a 40×/1.35 NA oil-immersion objective, a 405-nm laser, a 488-nm laser, a 561-nm laser, and a 640-nm laser, and controlled using NIS-Elements 4.51.00 software. The BFP, GFP, RFP, and near-infrared fluorescent signal were collected using 425–475 nm, 500–550 nm, 570–620 nm, and 663–738 nm filter, respectively.

### Live/Dead cell viability assay

The Live or Dead Cell Viability Assay kit (AAT Bioquest, 22789) was used to measure the viability of cells expressing PKU tags or other reported tags^25^. In brief, the cells were seeded in 24-well plates, transfected, and half of the cells were passaged every 24 hrs. On day 1 or day 7 post-transfection, the cells were detached using 0.25% trypsin-EDTA and collected in Eppendorf tubes. Non-transfected cells were fixed in 90% ethanol for 1 min at room temperature, and served as the negative control. After two washes with phosphate-buffered saline (PBS, pH 7.2), the cells were resuspended in 100 μl CytoCalcein Green/Propidium Iodide dye working solution and incubated at room temperature for 30 min, protected from light.

Flow cytometry was performed using a CytoFLEX S (Beckman) equipped with 405-nm, 488-nm, and 561-nm lasers. The following gating strategies were used to identify live and dead populations of cells expressing a certain protein. First, the acquired events were gated on FSC-A versus SSC-A to remove debris and aggregates, followed by gating for single cells on FSC-A versus FSC-H to exclude doublets. Second, cells expressing the expected tags were gated by the PB450 (tagBFP). Third, a dead cell gate was set on propidium iodide (PE channel)-positive, Calcein (FITC channel)-negative cells; conversely, live cells were gated as Calcein-positive, propidium iodide–negative. During data acquisition, a minimum of 10,000 events were collected per sample. Data were analyzed using FlowJ v10.8.1 Software (BD Life Sciences), with all gates applied consistently across samples to ensure comparability. Cell viability was calculated by dividing the number of live cells by sum of the live cells and dead cells.

### Electrical stimulation and calcium imaging

Primary neurons were incubated in culture medium containing 5 μM Calbryte590 AM (ATT Bioquest) at 37°C for approximately 30 min prior to calcium imaging. Field electrical stimuli were delivered using a GRASS Instruments S88 stimulator (A-M Systems). Each stimulation burst consisted of 100 pulses (1 ms per pulse duration) delivered at 20 Hz for 5 s applied at 80 V. The interval between each burst was 5 min. Calcium imaging was performed using an A1R confocal microscope equipped with appropriate filters and detectors for measuring Calbryte590 and PKU-sfGFP fluorescence. The change in fluorescence intensity was analyzed using ImageJ software (NIH). Regions of interest (ROIs) were selected in order to quantify the calcium in individual cells. The change in fluorescence (ΔR/R_0_) was calculated using the formula [(R−R_0_)/R_0_], where R is the fluorescent intensity ratio of Calbryte590 to PKU-sfGFP, and R_0_ is the baseline ratio. Statistical analyses were performed using OriginPro 2020b (OriginLab). Traces and summary graphs were generated using OriginPro 2020b (OriginLab).

### PKU tag assembly efficiency analysis

Imaging data were processed using ImageJ (1.54f) software (NIH). Cells expressing PKU tags were imaged using an Operetta system equipped with a 60× objective, generating *z*-stack images with a step size of 0.5 µm to create maximum projection images. The first filtering step involved calculating the average fluorescence intensity ratio between the cell and the background (i.e., an area without cells). Cells with a ratio < 1.3 were excluded from quantification due to their very low expression levels. Cells in which the ratio was ≥ 1.3 were identified as PKU tag-expressing. Next, a line was drawn through the brightest region of the assembled area as observed visually. The ratio of the maximum fluorescence intensity of the assembled region to the minimum value (background within the cell) along this line was then calculated. If this ratio was ≥ 1.4, the cell was classified as “assembled”, indicating that the PKU tag showed an assembled pattern that was clearly distinguishable from the background. If the ratio was < 1.4, the cell was classified as “non-assembled,” as the assembled pattern did not show a clear contrast compared to the background. Based on these criteria, we used assembled and non-assembled cells to quantify the assembly efficiency of each PKU tag.

### Double-blind recognition test

A total of nine participants were recruited, and seven different groups of PKU tags (three spherical tags and four filamentous tags) were used for double-blind recognition test. For each type of tag, images were collected and randomly divided into training (n=8 images) and test (n=35 images) sets. During the learning phase, the participants were instructed to learn the patterns of tags from the different groups. Next, the participants were asked to identify and categorize the images in a shuffled test set, with the images randomly ordered. The accuracy of participants’ classification was calculated on a scale from 0 to 1, with 1 indicating perfect recognition and the complete differentiation of all tags. A heatmap was then generated using OriginPro 2020b (OriginLab).

### Viral injections

Mice were anesthetized with an intraperitoneal injection of tribromoethanol (Avertin; 500 mg/kg body weight), and placed on a stereotaxic frame (RWD Life Science).

For the experiments in **Fig. 3b-c**, a total volume of 300 nl of mixed virus was injected into the primary motor cortex (M1; AP: +0.74 mm relative to Bregma; ML: ±1.0 mm relative to Bregma; DV: −0.65 mm from the dura) of wild-type female C57BL/6J mice (6–9 weeks of age) at a rate of 50 nl/min using a glass pipette and a micro-syringe pump (Nanoliter 2000 Injector, World Precision Instruments). The following mixtures of viruses was injected: AAV9-CaMKII-S1-tagBFP and AAV9-hSyn-EGFP (**Fig. 3b1**); AAV9-EFS-DIO-F1-mKate2 and AAV9-hSyn-EGFP-P2A-Cre (**Fig. 3b2**); AAV9-CaMKII-F5-tagBFP-nls and AAV9-hSyn-EGFP (**Fig. 3b4**); and AAV9-CaMKII-S1-tagBFP, AAV9-hSyn-F1-sfGFP and hSyn-tdTomato (**Fig. 3c**). The animals were sacrificed 3−4 weeks after injection.

For the experiments in **Fig. 3d**, a total volume of 300 nl of mixed viruses (AAV9-EFS-DIO-F1-mKate2, AAV9-hSyn-EGFP, and either AAV9-CaMKII-S1-tagBFP or AAV9-CaMKII-S1-mKate2) was injected into either M1 region (using the coordinates described above) or the primary visual cortex (V1; AP: −3.9 mm relative to Bregma; ML: ±2.6 mm relative to Bregma; DV: −0.5 mm from the dura) of VIP-Cre mice (6–9 weeks of age). The animals were sacrificed 3–4 weeks after injection.

For the experiments in **Fig. 3e** and **Extended Fig. 3**, AAV9-hSyn-S1-sfGFP, AAV9-hSyn-F1-sfGFP, AAV9-hSyn-S6-sfGFP-nls, AAV9-hSyn-F5-sfGFP-nls, AAV9-hSyn-sfGFP, or AAV9-hSyn-sfGFP-nls virus was injected into the basolateral amygdala (BLA; AP: −1.4 mm relative to Bregma; ML: −3 mm relative to Bregma; DV: −4.5 mm from the dura); 1, 3, 5, and 9 weeks after virus injection, the mice were sacrificed for imaging.

For the experiments in **Fig. 4b-c**, for anterograde tracing, two-step viral injections were performed as previously described^35^. In brief, 60 nl of AAV1-hSyn-Cre-P2A-GFP was injected into V1 using the coordinates described above. After 2–7 days, a second injection of AAV9-EFS-DIO-F1-mKate2 or AAV9-EFS-DIO-S1-tagBFP was administered (60 nl per site) into the ipsilateral superior colliculus (SC; AP: −3.9 mm relative to Bregma; ML: ±0.8 mm relative to Bregma; DV: −1.5 mm from the dura). The animals were typically euthanized 4 weeks after injection to allow sufficient time for viral transport and transgene expression.

For the experiments in **Fig. 4d**, 90 nl of AAV2-retro-hSyn-Flp-EGFP was injected into the SC using the coordinates described above. After 2–7 days, a second injection of mixed virus (AAV9-EFS-fDIO-S1-tagBFP and AAV9-EF1a-fDIO-GFP; 90 nl) was injected into the ipsilateral V1 using coordinates described above. The animals were sacrificed 4 weeks after injection.

For the experiments in **Fig. 4e**, 90 nl of AAV2-retro-hSyn-sfGFP-P2A-nls-Dre was injected into the lateral posterior (LP; AP: −2.3 mm relative to Bregma; ML: ±1.5 mm relative to Bregma; DV: −2.3mm from the dura). After 2–7 days, a second injection of AAV9-EFS-dDIO-F1-tagBFP (60 nl) was injected into the ipsilateral SC using the coordinates described above. The animals were sacrificed 4 weeks after injection.

For the experiments in **Fig. 6d**, a total volume of 300 nl of mixed viruses (either AAV9-hSyn-S1-sfGFP-APEX2 or AAV9-hSyn-F1-sfGFP-APEX2 and AAV9-hSyn-tdTomato) was injected into the M1 region of wild-type mice. The animals were sacrificed 3−4 weeks after injection.

For the experiments in **Fig. 6e**, a total volume of 300 nl of AAV9-EFS-DIO-F1-tagBFP-APEX2 was injected into the M1 region of Gad2-Cre mice. One month later, an additional 300 nl of AAV9-CaMKII-S1-sfGFP-APEX2 was injected into the same region. The animals were sacrificed 3–4 weeks after the secondary injection.

### *In utero* electroporation (IUE)

For the experiments in **Fig. 3b3**, a plasmid mixture containing EFS-S6-sfGFP-nls and CMV-mCherry was transfected into L2/3 pyramidal neurons in the barrel cortex by IUE. In brief, embryonic day 14.5 (E14.5) timed-pregnant female C57BL/6J mice were deeply anesthetized with pentobarbital (80 mg/kg body weight). The uterine horns were exposed and periodically rinsed with warm sterile PBS. A 1–2 μl volume of the plasmid DNA mixture (1–2 μg/μl) was injected into the lateral ventricle of the embryo *in utero* via a glass electrode. Fast Green FCF dye (Millipore Sigma) was included in the DNA mixture in order to visualize the mixture during injection. Five square-wave current electric pulses (34 V, 50-ms duration, 1 Hz) were delivered twice using 5-mm round plate electrodes (ECM830, BTX). The region of transfection was adjusted by slightly changing the orientation and position of the electrodes. After electroporation and surgery, the mice were kept warm with a heating pad until they recovered from anesthesia.

### Brain dissection and sectioning

After euthanasia, the mice were perfused with PBS followed by a 4% (w/v) paraformaldehyde (PFA, Sigma) solution. Next, the brains were dissected and incubated in 4% PFA at 4°C overnight. The fixed brains were sectioned into 40-μm-thick coronal slices in PBS with a VT1200 vibratome (Leica). The brain slices were stored in PBS at 4°C until use.

### Fluorescence imaging of mouse brain slices

For the experiments in **Fig. 3**, **Fig.4** and **Fig. 6**, brain slices containing the corresponding brain regions were transferred to a 24-well plate. Depending on the color of the expressed protein, the cell nuclei were stained with either DAPI (Beyotime) or DRAQ7 (BioLegend). Then slices were rinsed in PBS and mounted onto glass slides. To preserve fluorescence, the slides were stored in the dark at 4°C until imaging.

Brain slices were imaged using a Ti-E A1R inverted confocal microscope (Nikon) at various *z*-planes, and maximum intensity projection was performed to generate the final images.

### Transcriptome-wide RNA-seq analysis

Female C57BL/6J mice (6 weeks of age) were anesthetized with an intraperitoneal injection of tribromoethanol (Avertin; 500 mg/kg body weight). A 300 nl volume of AAV9-hSyn-S1-sfGFP (1 × 10^12), AAV9-hSyn-F1-sfGFP (2.5 × 10^13 vg/ml), AAV9-hSyn-S6-sfGFP-nls (4 × 10^12 vg/ml), AAV9-hSyn-F5-sfGFP-nls (4 × 10^12 vg/ml), AAV9-hSyn-sfGFP (1 × 10^12 or 2.5 × 10^13 vg/ml), or AAV9-hSyn-sfGFP-nls (4 × 10^12 vg/ml) virus was bilaterally injected into the M1 region (AP: +1.0 mm relative to Bregma; ML: ±1 mm relative to Bregma; DV: −0.65 mm from the dura). Three weeks after injection, the M1 region was dissected and snap-frozen in liquid nitrogen for RNA extraction. mRNA libraries were constructed and sequenced by AZENTA Life Sciences using Illumina HiSeq/ Illumina Novaseq/MGI2000 instrument. Sequencing data were filtered using Cutadapt V1.9.1 (Phred cutoff: 20; error rate: 0.1; adapter overlap: 1bp; min. length: 75; proportion of N: 0.1). The data were aligned to the reference genome using Hisat2 v2.0.1, and gene expression levels were estimated from the paired-end clean data using HTSeq v0.6.1. Heatmaps of FPKM (fragments per kilobase of transcript per million mapped reads) for selected immune-related genes were generated using R v4.2.2.

### Behavior assays of mice

Female C57BL/6J mice (8 weeks of age) were anesthetized, and AAV9-hSyn-S1-sfGFP, AAV9-hSyn-F1-sfGFP, AAV9-hSyn-S6-sfGFP-nls, AAV9-hSyn-F5-sfGFP-nls, AAV9-hSyn-sfGFP, or AAV9-hSyn-sfGFP-nls was injected into the BLA (AP: −1.4 mm relative to Bregma; ML: −3 mm relative to Bregma; DV: −4.5 mm from the dura). Three weeks and two months after virus injection, the mice were subjected to various behavioral tests in the following order and light intensities: open field test (13 lux), elevated plus maze test (330 lux), tail suspension test (70 lux), and forced swimming test (70 lux). During the behavioral tests, the mice were housed individually. The behavior during the open field test was analyzed using Activity Monitor 7 software (Med Associates), and all other tests were analyzed using Smart 3.0 (Panlab) software.

#### Open field test

The open field test was performed in an ENV-510 test chamber (27.3 × 27.3 × 20.3 cm, Med Associates). The chamber was equipped with infrared photo beams, which were used to evaluate spontaneous mouse locomotor activity. At the start of the test, the mouse was placed in the center of the arena and allowed to freely explore the environment for 10 min. The chambers were thoroughly cleaned with 75% ethanol between subjects. Mouse movement was tracked and analyzed for the time spent in the central zone (14.29 × 14.29 cm), the number of entries into the central zone, and the total distance traveled.

#### Elevated plus maze test

The elevated plus maze consists of two opposing open arms (30 × 5 cm) without walls and two opposing closed arms (30 × 5 × 15 cm) with opaque walls, with these four arms connected by a central platform (5 × 5 cm). The maze is elevated 50 cm above the floor. At the beginning of each session, the mouse was placed in the center platform facing one of the open arms and allowed to explore the maze for 5 min. Locomotion trajectories were recorded using a video camera. The arena was thoroughly cleaned with 75% ethanol between subjects. The time and number of entries into the various arms were quantified and analyzed.

#### Tail suspension test

The mouse was suspended by the tail approximately 40 cm above the floor, preventing any other contact or climbing during the assay. A 6-min session was then recorded, and the total time spent immobile during a 2−6-min period was analyzed.

#### Forced swim test

The mouse was placed in a transparent cylinder (12 cm in diameter and 30 cm in height), filled with water to a depth of 15 cm and maintained at 23–24°C. After each 6-min session, the mouse was dried with paper towels and returned to its home cage. The total immobility time during a 2−6-min period was recorded, and the mouse was considered immobile if it did not make any active movements.

### DAB staining and imaging of cultured cells

This procedure was performed as reported previously, with slight modification^15, 18^. In brief, cultured cells expressing the APEX2-related constructs were fixed using 2% glutaraldehyde (GA, Sigma) in 0.1 M phosphate buffer (PB, pH 7.2–7.4) at room temperature, then quickly moved to ice and incubated for 45–60 min. The cells were rinsed three times with chilled 0.1 M PB, for 10 min each. After rinsing, a solution containing 0.5 mg/mL 3,3-diaminobenzidine (DAB, D5905, Sigma) and 0.03% (v/v) H_2_O_2_ in 0.1 M PB was added to the cells. The reaction was monitored by the appearance of a dark brown color in the cells and stopped by replacing the DAB solution with 0.1 M PB. The samples were rinsed three times with 0.1 M PB at 4°C and then imaged using a Virtual Slide System (VS120-S6-W) brightfield microscope.

### DAB staining and imaging of brain slices

This procedure was performed as reported previously, with slight modifications^18, 19^. The mice were anesthetized and transcardially perfused with 0.1 M PB to remove the blood, followed by pre-fixation with 1% GA and 4% PFA in 0.1 M PB. The brains were dissected and post-fixed in 2.5% GA in 0.1 M PB at 4°C for at least 2 hrs (or overnight if needed). Following fixation, the brains were sectioned into approximately 100–200-μm-thick slices containing the motor cortex using a Leica VT1200 vibratome. The brain slices were then rinsed three times with 0.1 M PB at 4°C, for 10 min each. Following pre-incubation in filtered 0.3 mg/mL DAB in 0.1 M PB in the dark for 30 min, 0.03% (v/v) H_2_O_2_ (final concentration) was added to the samples for APEX2 oxidation. The reaction was monitored by the appearance of a dark brown color in the tissues and stopped by replacing the DAB solution with 0.1 M PB. The samples were rinsed three times with 0.1 M PB at 4°C and then imaged using a Virtual Slide System (VS120-S6-W) brightfield microscope.

### EM sample preparation of cultured cells

The initial steps of fixation and DAB staining of cultured cells were performed as described above. After these steps, 2% osmium tetroxide (OsO₄) in 0.1 M PB was added to each sample and incubated for 1 hr in the dark at room temperature. The samples were rinsed three times with ddH_2_O at 4°C, followed by overnight incubation in 1% aqueous uranyl acetate (UA) at 4°C in the dark. The samples were rinsed again with ddH_2_O three times for 10 min each at 4°C. The samples were then dehydrated using a graded ethanol series (30%, 50%, 75%, 85%, 95%, and 100%) at 4°C, followed by two rinses in 100% ethanol at room temperature, for 10 min each. Infiltration was carried out using a 1:3, 1:1, and 3:1 ratio of Epon 812 resin (Electron Microscopy Sciences) to 100% ethanol for 2 hrs each, followed by three infiltrations with freshly made pure Epon 812 for 1 hr each at room temperature. Finally, the samples were embedded in fresh resin and placed in a vacuum oven at 65°C for 24 hrs for polymerization. The samples were sectioned using a Leica EM UC7 ultramicrotome with Diatome diamond knives, and ultra-thin sections (80 nm) were picked up on slot grids.

### EM sample preparation of brain slices

The initial steps of fixation and DAB staining of brain slices were performed as described above. After these steps, the DAB-positive region of the motor cortex was excised from the brain slices. The excised tissue was incubated in 2% osmium tetroxide (OsO₄) in 0.1 M PB for 1 hr in the dark at room temperature, then washed with ddH_2_O three times for 10 min each at 4°C. The tissue was incubated overnight in 1% aqueous UA at 4°C in the dark, then washed three times with ddH_2_O for 10 min each at 4°C. The tissues were then dehydrated using a graded ethanol series (30%, 50%, 75%, 85%, 95%, and 100%) at 4°C, followed by two rinses in 100% ethanol and one rinse in acetone at room temperature, for 10 min each. Infiltration was carried out using a 1:3, 1:1, and 3:1 ratio of Epon 812 resin to acetone for 2 hrs each at room temperature. The samples were then processed as described previously.

### Transmission EM

TEM images were acquired at 80 kV accelerating voltage using a JEM-1400 transmission electron microscope (JEOL, Japan) equipped with a XAROSA camera (Emsis GmbH, Muenster, Germany). Images were saved in.tif format, and the darkness intensity of regions expressing PKU-sfGFP-APEX2 was quantified using ImageJ software (NIH).

### FIB-SEM

For correlative light and electron microscopy (CLEM) experiments, EM samples prepared as described above were imaged using FIB-SEM (FEI Helios Nanolab G3 UC). First, sample blocks were affixed to sample stubs using conductive adhesive tabs. All surfaces of the block—except for the sample itself—were coated with colloidal silver paint and dried in a vacuum chamber. The samples were then sputter-coated with palladium for 120 s. A 2.5-nA gallium beam was used to mill the samples at 10 nm per layer. Following each milling step, the process was paused to image the freshly exposed surface using a 2-kV acceleration potential using the in-column energy-selective backscattered electron detector (ICD). The pixel dwell time for imaging was set to 3 µs. This sequence of milling and imaging was repeated in an automated manner, requiring minimal supervision, and typically producing several hundred images. The resolution of the images in the *XY* plane was set to approximately 6.30 nm/pixel for F1-sfGFP-APEX2, 13.02 nm/pixel for S7-sfGFP-APEX2-nls, and 16.28 nm/pixel for F5-sfGFP-APEX2-nls. The resolution in the *Z*-axis, determined by the thickness of the removed layers, was 10 nm for all samples. Image dimensions were 20.13 μm (width) × 25.81 μm (height) × 5.62 μm (milling length) for F1-sfGFP-APEX2, 39.99 μm (width) × 26.65 μm (height) × 7.16 nm (milling length) for S7-sfGFP-APEX2-nls, and 49.98 μm (width) × 33.32 μm (height) × 7.2 μm (milling length) for F5-sfGFP-APEX2-nls. The acquired image stacks were aligned, and the electron-dense labels were reconstructed using Amira 2022.1 software.

### Immunostaining of Qt tag-related samples

HEK cells were plated on 12-mm glass coverslips and transfected with either EFS-mScarlet-IP2A-1M-Qt-flag-nls or EFS-mScarlet-IP2A-2M-Qt-flag. After 36 hrs, cells were fixed in 4% PFA in PBS for 15 min, followed by three washes with PBS, each lasting 10 min. The cells were then permeabilized with 0.2% Triton X-100 in PBS for 20 min and washed three more times with PBS for 10 min each. After that, cells were blocked in 5% bovine serum albumin (BSA) in PBS for 1 hr at room temperature and then incubated overnight at 4 °C with Anti-Flag primary antibodies (#8146, Cell Signaling). After incubation, the cells were washed three times with PBS for 10 min each. Secondary antibodies (goat anti-mouse Alexa Fluor 405) were then added, and incubated at room temperature for 1 hr, followed by three washes with PBS for 10 min each. Finally, the cells were stained with DRAQ7 (catalog no. 424001, BioLegend) for 1 hr at room temperature, mounted on slides, and imaged using confocal microscopy as previously described.

### EM preparation of Qt tag-related samples

This EM procedure was performed as described previously, with minor modifications^25^. The main differences in the protocol include the exclusion of propylene oxide for dehydration due to its corrosive effect on the dish, and the use of our accessible Epon 812 for embedding. These two steps are unrelated to heavy metal staining and are not expected to impact the electron-dense labeling.

HEK cells were plated on the dishes (MATTEK, P35G-1.5-14-C-GRD) and transfected with either EFS-mScarlet-IP2A-1M-Qt-flag-nls or EFS-mScarlet-IP2A-2M-Qt-flag.

After 36 hrs, cells were imaged using an A1 confocal microscope and then fixed with 2.5% GA in 0.1 M sodium cacodylate buffer (pH 6.7–7.0) for 30 min. After removal of the fixative, the cells were postfixed with a 1:1 mixture of 4% OsO_4_ and 0.3 M sodium cacodylate buffer, containing 3% potassium hexacyanoferrate (II) for 20 min on ice. The postfixative solution was removed, and the samples were washed twice with 0.1 M sodium cacodylate buffer for 2 min, followed by 4 × 10 min ddH₂O washing steps. Next, the samples were stained with 1% tannic acid for 10 min at room temperature, then decanted and washed five times with ddH₂O for 5 min each. Subsequently, the samples were treated with 1% UA solution for 30 min at room temperature, followed by 5 × 5 min washes with ddH₂O. The samples were then treated with lead aspartate (containing 0.02 M lead nitrate and 0.03 M aspartic acid, Sigma-Aldrich) for 15 min at 60°C, followed by 5 × 8 min washes with ddH₂O.

After washing, the samples were then dehydrated using a graded ethanol series (30%, 50%, 75%, 85%, 95%, and 100%) at 4°C, followed by two rinses in 100% ethanol at room temperature, for 10 min each. The ethanol was discarded, and the samples were subjected to three infiltrations with a 1:3, 1:1, and 3:1 ratio of Epon 812 resin to 100% ethanol for 2 hrs each. This was followed by three infiltrations with freshly prepared pure Epon 812 resin for 1 hr each at room temperature. Finally, the samples were embedded in fresh resin and polymerized in a vacuum oven at 65°C for 24 hrs. The samples were then sectioned using a Leica EM UC7 ultramicrotome (70 nm). EM images were acquired at 80 kV accelerating voltage using a JEM-1400. Images used for darkness quantification were captured at a magnification of 25,000x.

### Quantification and statistical analysis

Cartoons and schematic diagrams were created using BioRender (http://www.biorender.com). Data were plotted using OriginPro 2021b (OriginLab). Except where indicated otherwise, all summary data are reported as the mean ± s.e.m. All data were assumed to be distributed normally, and equal variances were formally tested. Differences were analyzed using one-way ANOVA with Tukey’s multiple comparison test. In all figures, *P < 0.05, **P < 0.01, ***P < 0.001, and NS, not significant (P ≥ 0.05).

**Extended Data Fig. 1.**
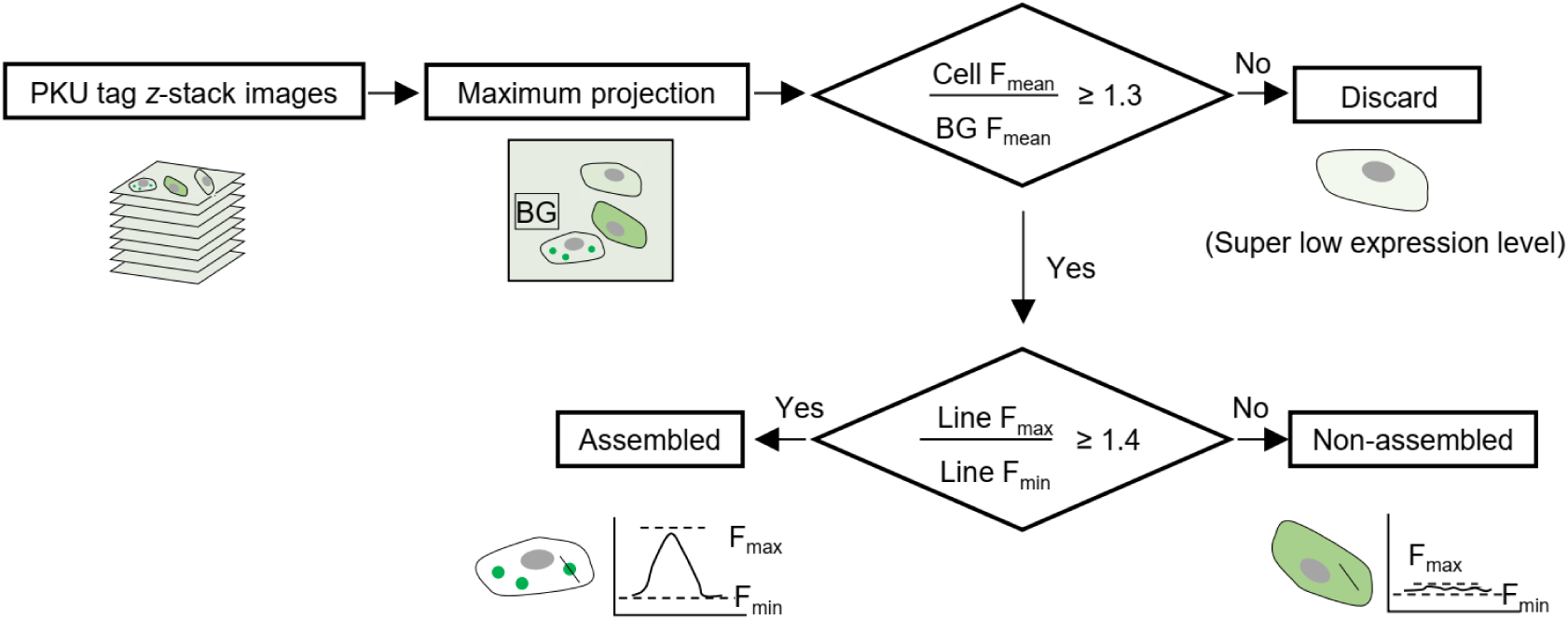
| Criteria for defining “assembled” and “non-assembled” cells. Cells expressing PKU tags were imaged across different *z*-planes to generate maximum projection images. The cells were then classified as either “assembled” or “non-assembled” based on the following two criteria: *i*) the ratio of the cell’s mean fluorescence intensity to the background fluorescence intensity (i.e., area without cells) and *ii*) the ratio of the assembled region’s maximum fluorescence intensity to the cell’s cytosolic background. These two ratios were then used to quantify the assembly efficiency of each PKU tag.

**Extended Data Fig. 2.**
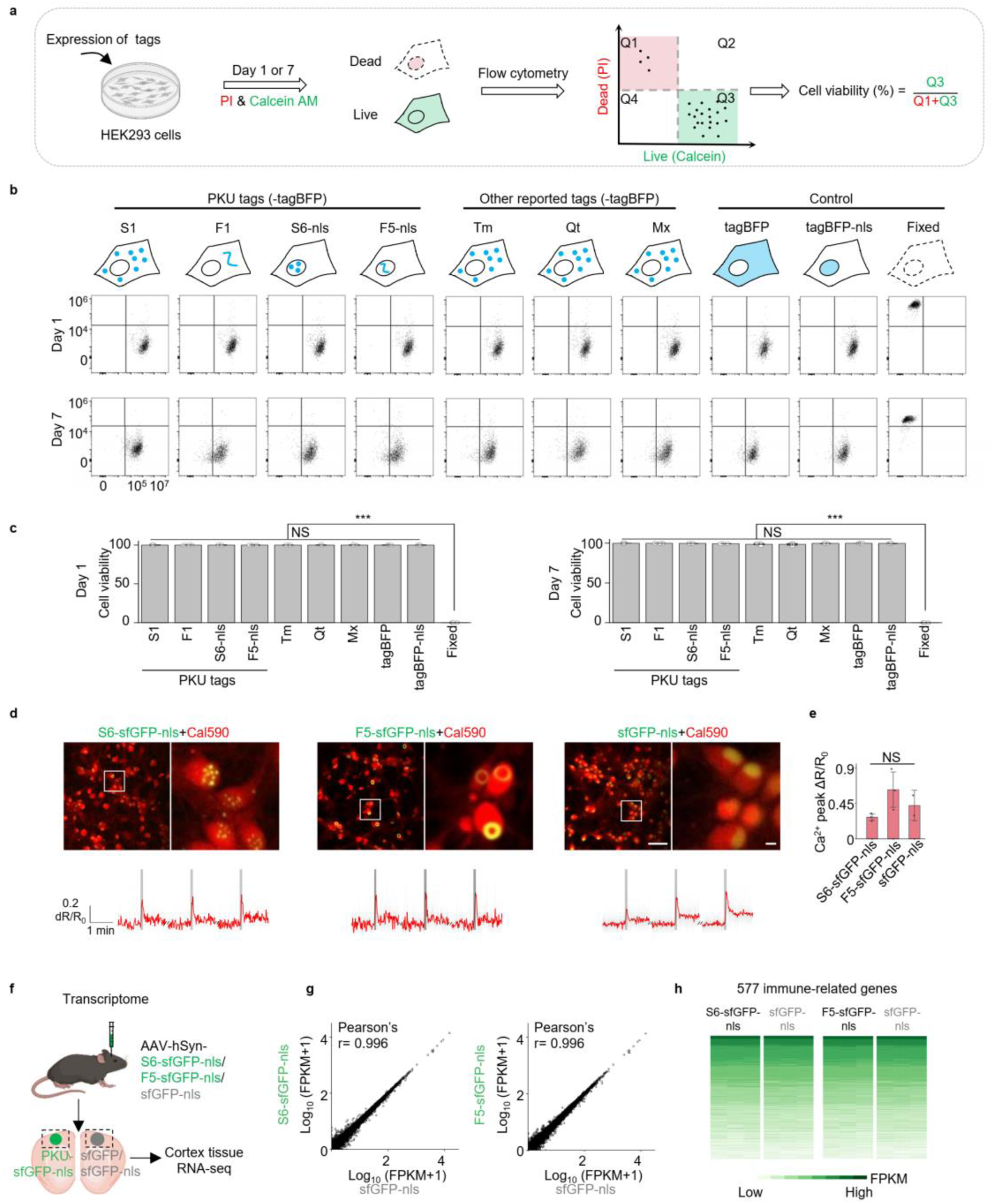
| Expression of a PKU tag has no measurable effect on cell viability, physiology or transcriptome. **a**. Schematic diagram depicting the protocol for measuring cell viability and the formula for calculating cell viability. **b.** Representative flow cytometry plots of cells expressing the indicated PKU tags fused with tagBFP for 1 or 7 days. Tm, Qt and Mx are previously reported self-assembled barcodes^25^, also fused with tagBFP. Cells expressing cytosolic tagBFP or nuclear-localized tagBFP (tagBFP-nls) served as positive controls, while fixed non-transfected cells served as a negative control. **c.** Summary of cell viability in cells expressing the indicated tags, measured 1 and 7 days after transfection. **d-e**. Expressing PKU tags in cultured rat hippocampal neurons does not affect their cellular physiology **d**, Representative fluorescent images (top) and average calcium response traces (bottom) measured in cultured rat hippocampal neurons expressing S6-sfGFP-nls (left), F5-sfGFP-nls (middle), or sfGFP-nls (right); where indicated by the dark gray shaded area, trains of electrical stimuli (100 pulses delivered at 20 Hz, 1 ms per pulse duration) were applied. Calcium response signals (ΔR/R_0_) were normalized to sfGFP. The red trace represents the mean ΔR/R_0_ across neurons in the same coverslip (n=47 neurons). The light gray traces denote individual neuronal calcium responses. The white boxes show the magnified regions. **e**. Summary of peak (ΔR/R_0_) measured in neurons expressing S6-sfGFP-nls (n=3 coverslips), F5-sfGFP-nls (n=3 coverslips) or sfGFP-nls (n=2 coverslips) in response to electrical stimulation. **f-h**. RNA-seq analysis shows that expressing PKU tags does not affect the cellular transcriptome. **f**. Schematic diagram depicting the experimental strategy in which PKU-sfGFP-nls and sfGFP-nls were expressed in the mouse motor cortex for 3 weeks, after which the corresponding tissues were collected for RNA-seq analysis. **g**. Pairwise comparisons of transcriptomes between motor cortex samples expressing S6-sfGFP-nls vs. sfGFP-nls (left), and between F5-sfGFP-nls vs. sfGFP-nls (right) (n=3 mice per group). The Pearson′s correlation coefficients are shown. **h**. Expression of selected immune-related genes was compared between motor cortex samples expressing PKU-sfGFP-nls vs. sfGFP-nls (n=3 mice per group).

**Extended Data Fig. 3.**
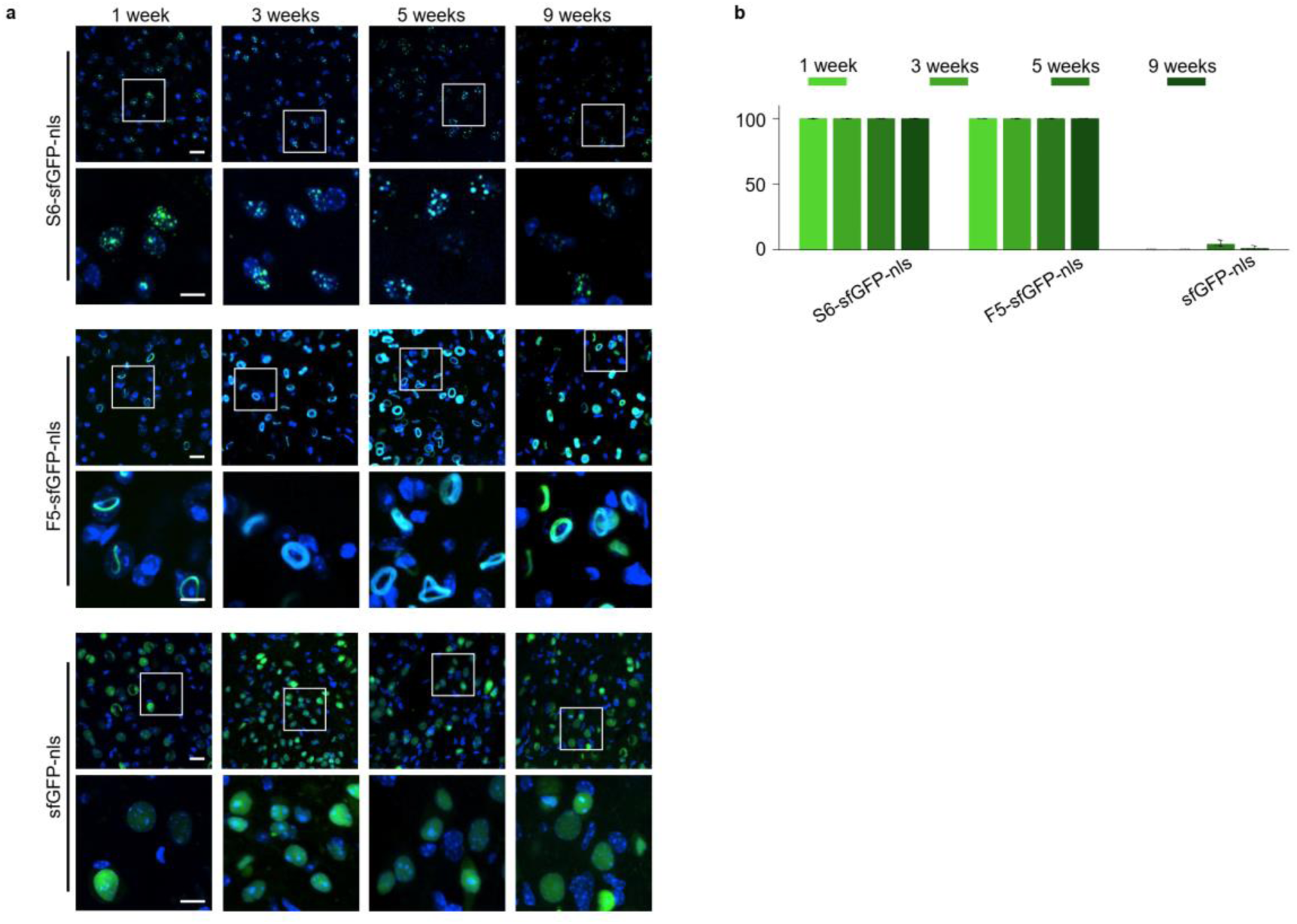
| PKU tags expressed *in vivo* maintained their assembly patterns for up to 9 weeks. **a**. Representative images of sections expressing the indicated PKU tags 1, 3, 5 and 9 weeks after virus injection (top row), with magnified images of assembled patterns (bottom row). Scale bars: 20 μm (top) and 10 μm (bottom). **b.** Summary of the assembly efficiency of S6-sfGFP-nls, F5-sfGFP-nls and sfGFP-nls 1, 3, 5, and 9 weeks after virus injection (n=3 brain slices per group).

**Extended Data Fig. 4.**
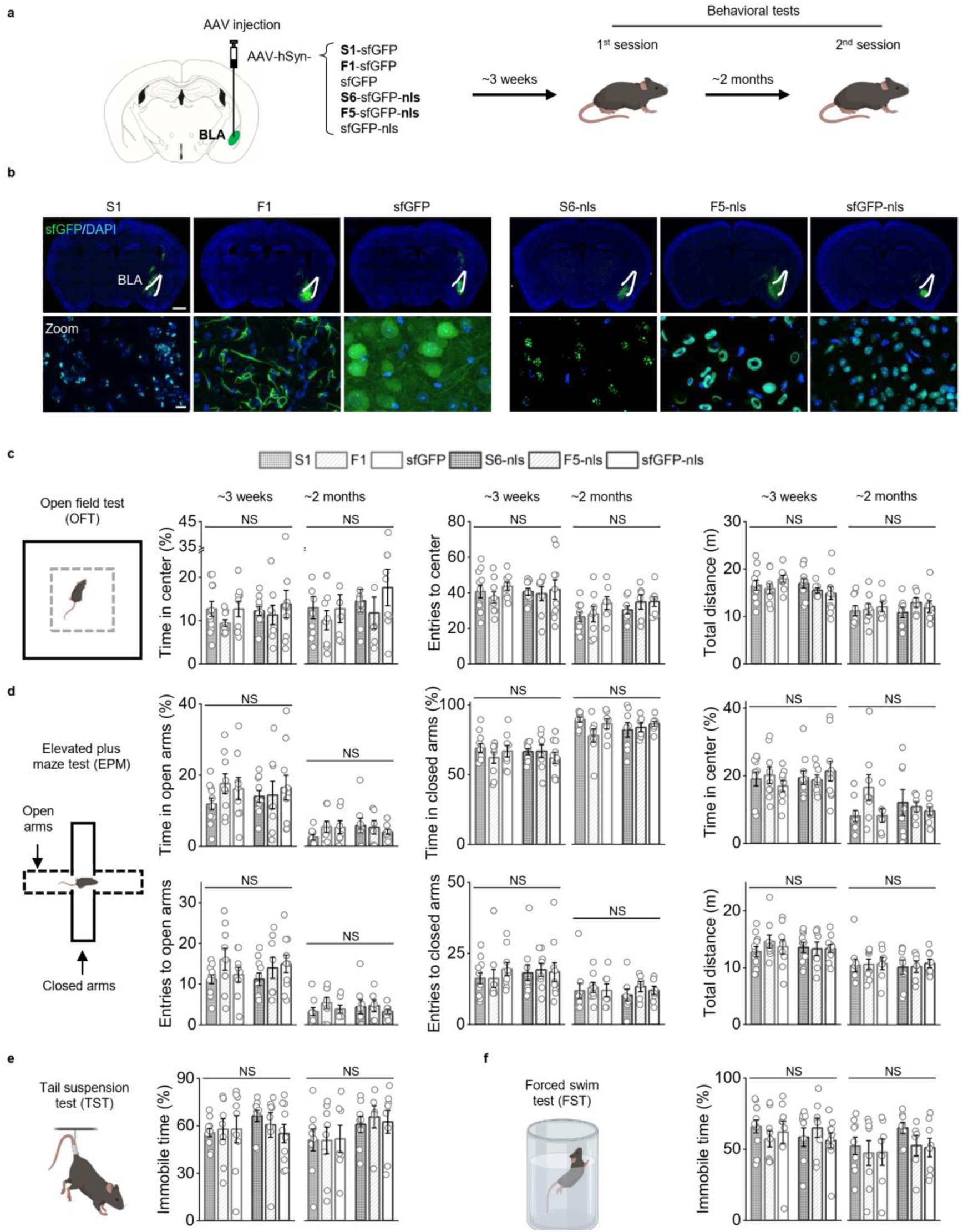
| Expressing PKU tags *in vivo* has no significant effect on animal behavior. **a**. Schematic diagram illustrating the experimental design. After expressing the indicated PKU tags via AAV, behavioral tests were performed at 3 weeks and 2 months. **b.** Representative fluorescence images of brain slices showing expression of the indicated PKU tags in the BLA (top) and magnified views showing the assembled patterns (bottom) 3 weeks after viral injection. The nuclei were stained with DAPI (blue). Scale bars: 1 mm (top) and 10 μm (bottom). **c–f**. Schematic depiction (left) and summary of results (right) measured in the open field test (**c**), elevated plus maze test (**d**), tail suspension test (**e**) and forced swim test (**f**) 3 weeks and 2 months after expressing the indicated PKU tags.

**Extended Data Fig. 5.**
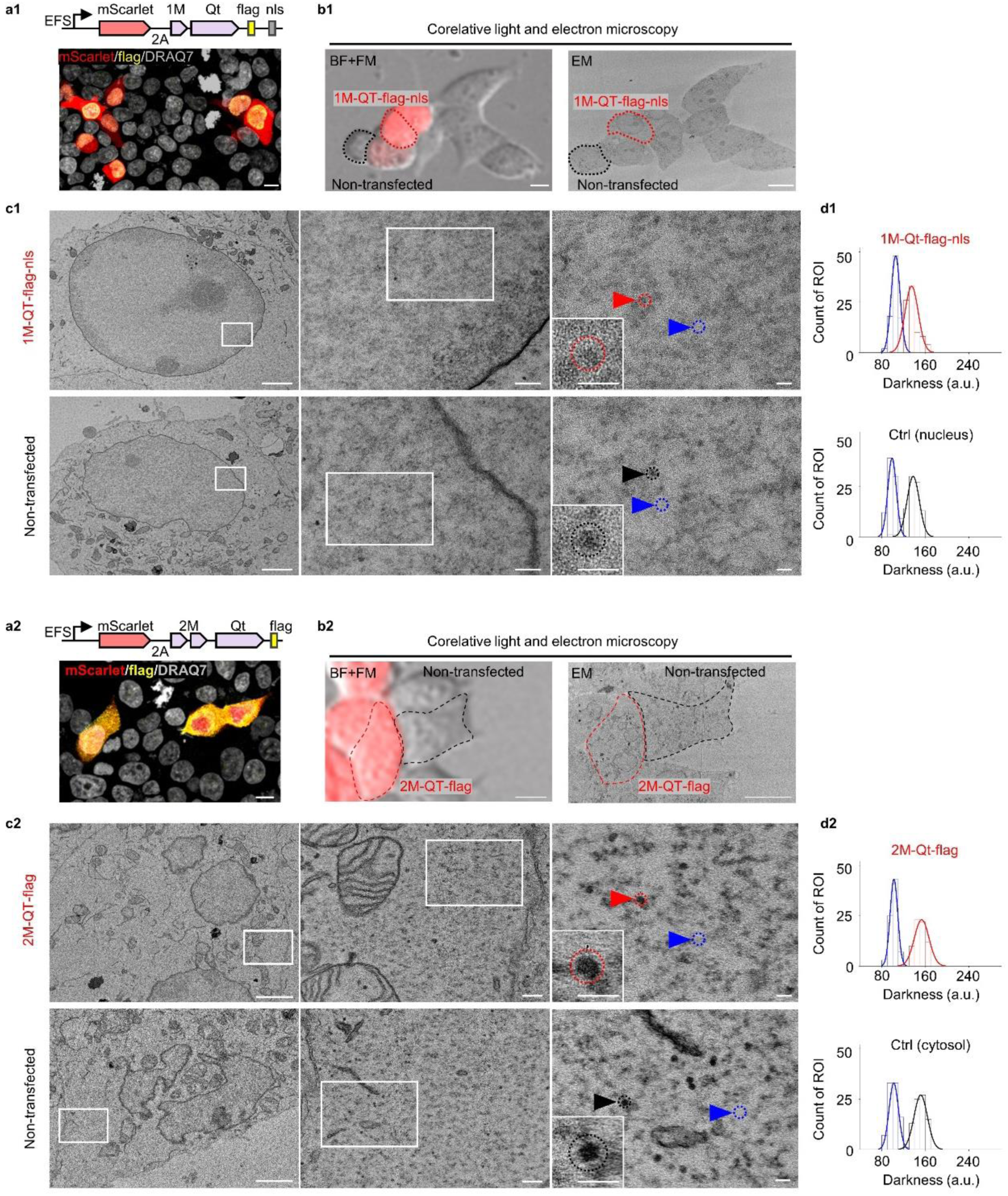
| Electron microscopy images of cells expressing reported Qt barcodes. **a**. Top: schematic diagram depicting the genetic constructs for expressing 1M-Qt-flag-nls (**a1**) and 2M-Qt-flag (**a2**) together with mScarlet in the HEK cells. Bottom: representative fluorescence images of the HEK cells expressing 1M-Qt-flag-nls (**a1**) and 2M-Qt-flag (**a2**). Flag epitopes are stained with antibody (yellow). Gray indicates nuclear staining with DRAQ7. Scale bars: 10 μm. **b.** Correlative light and electron microscopy images of cultured HEK cells transfected with 1M-Qt-flag-nls (**b1**) and 2M-Qt-flag (**b2**). Left: merged images of bright field (BF) and red fluorescence channel. Right: corresponding electron microscopy images of the same cells shown in the left panels. Red dashed lines indicate the outline of cells expressing 1M-Qt-flag-nls (**b1**) and 2M-Qt-flag (**b2**), whose EM images are shown in **c1** and **c2**. Black dashed lines indicate the non-transfected cells shown for comparison in **c1** and **c2**. Scale bars: 10 μm. **c.** Electron microscopy images of cells expressing 1M-Qt-flag-nls (**c1**, top) and 2M-Qt-flag (**c2**, top) and non-transfected cells (**c1** and **c2**, bottom). White boxes indicate the regions magnified in the right. Insets display increasingly magnified views. Red arrowheads highlight spherical electron-dense labeling potentially generated by 1M-Qt-nls (**c1**, red) and 2M-Qt (**c2**, red). Black arrowheads indicate other spherical electron-dense structures observed in the nucleus (**c1**, black) or cytosol (**c2**, black) of untransfected cells. Blue arrowheads mark nuclear (**c1**, blue) or cytosolic (**c2**, blue) background regions. The dashed circle outlines the region used for quantification of darkness distribution in **d**. Scale bars: 2 μm (left), 200 nm (middle), 50 nm (right), 50 nm (inset). **d.** Histogram of darkness distribution. Red represents the quantification of spherical electron-dense labeling potentially generated by 1M-Qt-nls (**c1**, red) and 2M-Qt (**c2**, red). Blue represents the quantification of nuclear background regions (**d1**, blue) and cytosolic background regions (**d2**, blue). Black represents the quantification of other spherical electron-dense labeling observed in the nuclei (**d1**, black) or cytosol (**d2**, black) of untransfected cells.

